# Mucosal host–microbe interactions associate with clinical phenotypes in inflammatory bowel disease

**DOI:** 10.1101/2022.06.04.494807

**Authors:** Shixian Hu, Arno R. Bourgonje, Ranko Gacesa, Bernadien H. Jansen, Johannes R. Björk, Amber Bangma, Iwan J. Hidding, Hendrik M. van Dullemen, Marijn C. Visschedijk, Klaas Nico Faber, Gerard Dijkstra, Hermie J. M. Harmsen, Eleonora A. M. Festen, Arnau Vich Vila, Lieke M. Spekhorst, Rinse K. Weersma

## Abstract

Dysregulation of gut mucosal host–microbe interactions is a central feature of inflammatory bowel disease (IBD). To study tissue-specific interactions, we performed transcriptomic (RNA-seq) and microbial (16S-rRNA-seq) profiling of 696 intestinal biopsies derived from 353 patients with IBD and controls. Analysis of transcript-bacteria interactions identified six distinct groups of inflammation-related pathways that were associated with intestinal microbiota, findings we could partially validate in an independent cohort. An increased abundance of *Bifidobacterium* was associated with higher expression of genes involved in fatty acid metabolism, while *Bacteroides* was associated with increased metallothionein signaling. In fibrostenotic Crohn’s disease, a transcriptional network dominated by immunoregulatory genes associated with *Lachnoclostridium* bacteria in non-stenotic tissue. In patients using TNF-α-antagonists, a transcriptional network dominated by fatty acid metabolism genes associated with *Ruminococcaceae*. Mucosal microbiota composition was associated with enrichment of specific intestinal cell types. Overall, we identify multiple host–microbe interactions that may guide microbiota-directed precision medicine.

## Main

Inflammatory bowel diseases (IBD), which encompass Crohn’s disease (CD) and ulcerative colitis (UC), are chronic inflammatory diseases of the gastrointestinal tract [1]. The pathogenesis of IBD is thought to be caused by a complex interplay between inherited and environmental factors, gut microbiota and the host immune system [2, 3]. Alterations in gut microbiota composition and functionality are commonly observed in patients with IBD, including decreased microbial diversity, decreased abundances of butyrate-producing bacteria and increased proportions of pathobionts [4–8].

Interactions between host genetics and the gut microbiome have been studied in both healthy subjects and patients with IBD. For example, we previously focused on host genome–gut microbiota interactions in the context of IBD [9]. However, in order to disentangle disease mechanisms that might underlie the etiology and progression of IBD, there should be a greater focus on mucosal gene expression studies [10].

Modulation of host mucosal gene expression by gut microbiota or effects of gene expression on microbial fitness may expose mechanisms that contribute to IBD pathogenesis, knowledge that could be utilized to explore novel therapeutic targets [11, 12]. Most studies, however, employ fecal sampling for microbiota characterization, which precludes analysis of local interactions and their immediate impact on host intestinal expression signatures. Such studies examining mucosal gene expression– microbiome associations in the context of IBD previously identified microbial groups associated with host transcripts from immune-mediated and inflammatory pathways [12–15]. In a longitudinal host–microbe interaction study, the chemokine genes *CXCL6* and *CCL20* were negatively associated with the relative abundances of *Eubacterium rectale* and *Streptococcus*, suggesting that these bacteria are more susceptible to the actions of these chemokines [13]. Another study found an inverse association between host expression of *DUOX2*, which produces reactive oxygen species (ROS), and the relative abundance of *Ruminococcaceae*, an association that may suggest ROS-mediated antibacterial effects [16]. However, few studies to date have been able to carry out comprehensive integrated analysis of IBD-associated interaction factors among mucosa-attached microbiota and host intestinal-gene expression.

Here we analyzed 696 fresh-frozen intestinal biopsies derived from 337 patients with IBD and 16 non-IBD controls for which we generated both mucosal transcriptomic and microbial characterization using bulk RNA-sequencing and 16S rRNA gene sequencing, respectively. We further combined both datasets to comprehensively investigate mutual mucosal host-microbe interactions and integrated these with the extensive clinical characteristics collected. Following this approach, we aimed to investigate mucosal host–microbe interactions while disentangling disease-, location- and inflammation-specific associations (a graphical representation of the study workflow is presented in **Figure 1**). Most importantly, we could study the associations between mucosal host– microbe interactions and clinical phenotypes of patients with IBD. Finally, we also sought to replicate our main results in data from a smaller independent, publicly available cohort [13].

**Figure 1.**
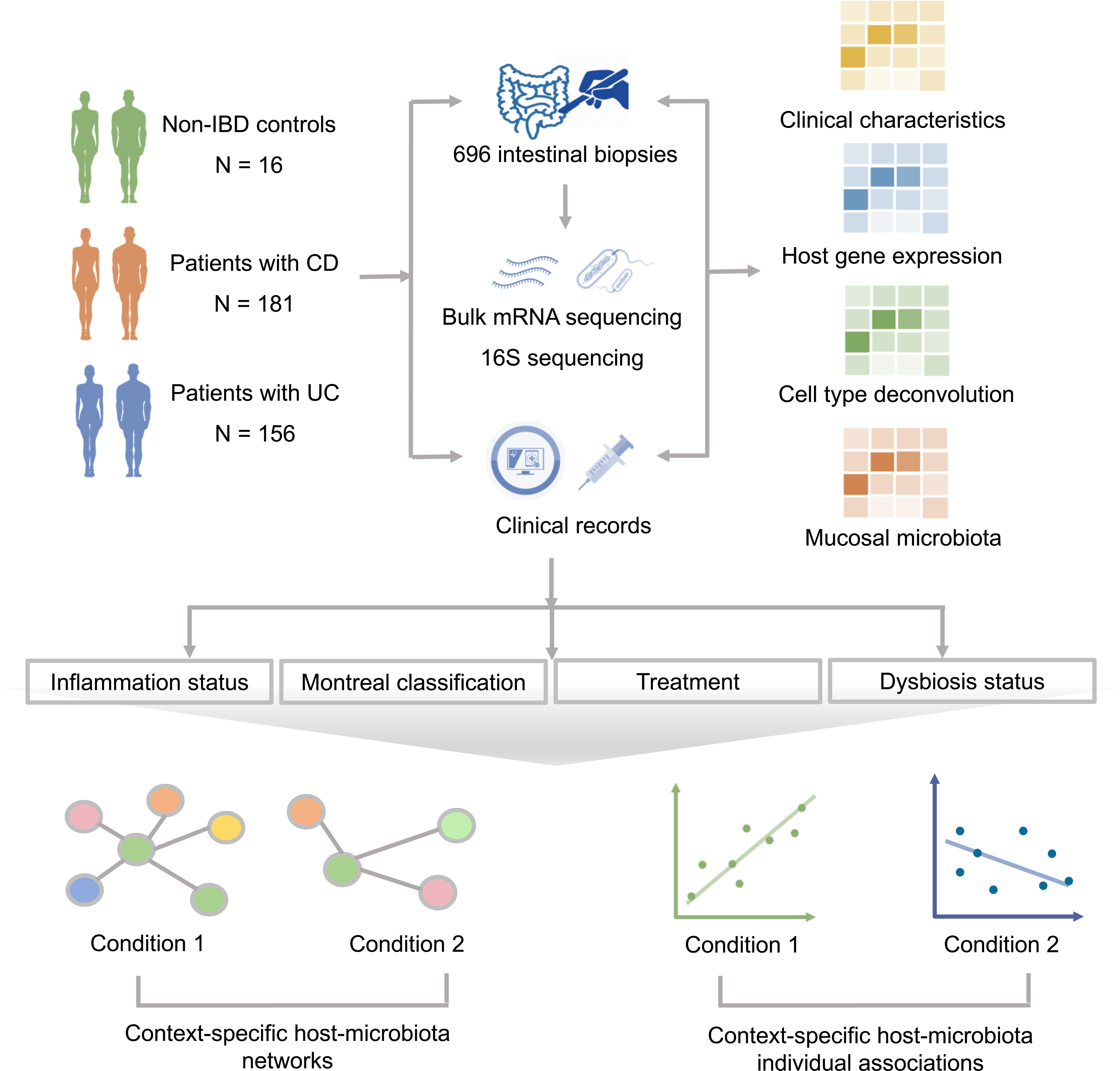
**Methodological workflow of the study**. The study cohort consisted of 337 patients with IBD (CD: *n*=181, UC: *n*=156) and 16 non-IBD controls, from whom 696 intestinal biopsies were collected (IBD: *n*=640, controls: *n*=56) and processed to perform bulk mucosal mRNA-sequencing and 16S gene rRNA sequencing. Detailed phenotypic data were extracted from clinical records for all study participants. In total, 251 ileal biopsies (CD: *n*=186, UC: *n*=56, controls: *n*=9) and 445 colonic biopsies (CD: *n*=165, UC: *n*=233, controls: *n*=47) were included: 212 biopsies derived from inflamed regions and 484 from non-inflamed regions. Mucosal gene expression and bacterial abundances were systematically analyzed in relation to different (clinical) phenotypes: presence of tissue inflammation, Montreal disease classification medication use (e.g. TNF-α-antagonists) and dysbiotic status. Pathway-based clustering and network analysis (Sparse-CCA and centrLCC analysis) and individual pairwise gene–taxa associations were investigated to identify host–microbiota interactions in different contexts. We then analyzed the degree to which mucosal microbiota could explain the variation in intestinal cell type–enrichment (estimated by deconvolution of bulk RNA-seq data). To confirm our main findings, we used publicly available mucosal 16S and RNA-seq datasets for external validation [13].

## Results

### Cohort description

Demographic and clinical characteristics of the study population are presented in **Table 1**. In total, we included 640 intestinal biopsies from 337 patients with IBD and 56 intestinal biopsies from 16 non-IBD controls. Biopsies were derived from the colon (64.4%) and ileum (35.6%), and patients with CD and UC were equally represented among inflamed (CD: 53.8%, UC: 46.2%) and non-inflamed (CD: 55.4%, UC: 44.6%) biopsies. Mean age and the proportion of smokers were higher among controls (*P*<0.01 and *P*=0.01, respectively). Among biopsies derived from patients with IBD, the proportion of steroid users was higher among patients from whom inflamed biopsies were collected (*P*<0.01). Remaining patient characteristics were evenly distributed among groups without significant differences.

**Table 1.** Demographic and clinical characteristics of the study population compared between the inflamed and non-inflamed dataset.

| Variable | Total | IBD |  | Non-IBD Controls | P-value |
| --- | --- | --- | --- | --- | --- |
|  |  | Inflamed biopsies | Non-inflamed biopsies |  |  |
|  | *n* = 696 | *n* = 212 | *n* = 428 | *n* = 56 |  |
| Biopsy inflammation, *n* (%) |  |  |  |  |  |
| *Inflamed* | 212 (30.5) | 212 (100) | - | - |  |
| *Non-inflamed* | 428 (61.5) | - | 428 (100) | - |  |
| Biopsy location, *n* (%) |  |  |  |  | <0.01 |
| *Ileum* | 248 (35.6) | 66 (31.1) | 173 (40.4) | 9 (16.1) |  |
| *Colon* | 448 (64.4) | 146 (68.9) | 255 (59.6) | 47 (83.9) |  |
| Diagnosis or control, *n* (%) |  |  |  |  | 0.74 |
| *CD* | 351 (50.4) | 114 (53.8) | 237 (55.4) | - |  |
| *UC* | 289 (41.5) | 98 (46.2) | 191 (44.6) | - |  |
| *Controls* | 56 (8.0) | - | - | 56 (100) |  |
| Age at biopsy (years) | 43.1 ± 15.3 | 43.0 ± 15.8 | 42.3 ± 15.5 | 45.5 ± 10.8 | <0.01 |
| Sex, *n* (%) |  |  |  |  |  |
| *Male* | 322 (46.3) | 92 (43.4) | 188 (43.9) | 42 (75.0) |  |
| *Female* | 374 (53.7) | 120 (56.6) | 240 (56.1) | 14 (25.0) |  |
| BMI (kg/m<sup>2</sup>) | 25.7 ± 4.5 | 25.7 ± 4.6 | 25.7 ± 4.7 | 24.7 ± 2.5 |  |
| Current smoking, *n* (%) |  |  |  |  | 0.01 |
| *Yes* | 145 (20.8) | 37 (17.5) | 88 (20.6) | 20 (35.7) |  |
| *No* | 551 (79.2) | 175 (82.5) | 340 (79.4) | 36 (64.3) |  |
| **Montreal classification** |  |  |  |  |  |
| Montreal Age (A), *n* (%) | 638 (99.7) | 212 (100) | 426 (99.5) | - | 0.95 |
| A1 (≤16 years) | 73 (11.4) | 23 (10.8) | 50 (11.7) | - |  |
| A2 (17–40 years) | 397 (62.0) | 132 (62.3) | 265 (62.2) | - |  |
| A3 (>40 years) | 168 (26.3) | 57 (26.9) | 111 (26.1) | - |  |
| Montreal Location (L), *n* (%) | 333 (94.9) | 108 (94.7) | 225 (94.9) | - | 0.83 |
| L1 (ileal disease) | 66 (19.8) | 18 (16.7) | 48 (21.3) | - |  |
| L2 (colonic disease) | 51 (15.3) | 17 (15.7) | 34 (15.1) | - |  |
| L3 (ileocolonic disease) | 172 (51.7) | 58 (53.7) | 114 (50.7) | - |  |
| L1 + L4 | 12 (3.6) | 3 (2.8) | 9 (4.0) | - |  |
| L2 + L4 | 6 (1.8) | 3 (2.8) | 3 (1.3) | - |  |
| L3 + L4 | 26 (7.8) | 9 (8.3) | 17 (7.6) | - |  |
| Montreal Behavior (B), *n* (%) | 333 (94.9) | 108 (94.7) | 225 (94.9) | - | 0.31 |
| B1 (non-stricturing, non-penetrating) | 146 (43.8) | 53 (49.1) | 93 (41.3) | - |  |
| B2 (stricturing) | 59 (17.7) | 20 (18.5) | 39 (17.3) | - |  |
| B3 (penetrating) | 30 (9.0) | 9 (8.3) | 21 (9.3) | - |  |
| B1 + P (perianal disease) | 35 (10.5) | 10 (9.3) | 25 (11.1) | - |  |
| B2 + P (perianal disease) | 48 (14.4) | 15 (13.9) | 33 (14.7) | - |  |
| B3 + P (perianal disease) | 15 (4.5) | 1 (0.9) | 14 (6.2) | - |  |
| Montreal Extension (E), *n* (%) | 246 (85.1) | 83 (84.7) | 163 (85.3) | - | 0.95 |
| E1 (proctitis) | 19 (7.7) | 7 (8.4) | 12 (7.4) | - |  |
| E2 (left-sided colitis) | 75 (30.5) | 25 (30.1) | 50 (30.7) | - |  |
| E3 (pancolitis) | 152 (61.8) | 51 (61.4) | 101 (62.0) | - |  |
| Montreal Severity (S), *n* (%) | 207 (71.6) | 68 (69.4) | 139 (72.8) |  | 0.70 |
| S0 (remission) | 11 (5.3) | 3 (4.4) | 8 (5.8) |  |  |
| S1 (mild) | 28 (13.5) | 9 (13.2) | 19 (13.7) |  |  |
| S2 (moderate) | 109 (52.7) | 33 (48.5) | 76 (54.7) |  |  |
| S3 (severe) | 59 (28.5) | 23 (33.8) | 36 (25.9) |  |  |
| **Medication use** |  |  |  |  |  |
| Aminosalicylates, *n* (%) | 271 (42.3) | 91 (42.9) | 180 (42.1) | - | 0.87 |
| Thiopurines, *n* (%) | 210 (32.8) | 66 (31.1) | 144 (33.6) | - | 0.53 |
| Steroids, *n* (%) | 262 (40.9) | 105 (49.5) | 157 (36.7) | - | <0.01 |
| Methotrexate, *n* (%) | 44 (6.9) | 18 (8.5) | 26 (6.1) | - | 0.32 |
| TNF-α-antagonists, *n* (%)<sup>†</sup> | 113 (17.7) | 35 (16.5) | 78 (18.4) | - | 0.58 |
| **Clinical disease activity** |  |  |  |  |  |
| HBI | 324 (92.3) | 104 (91.2) | 220 (92.8) | - | 0.18 |
| Remission (<5) | 205 (63.3) | 60 (57.7) | 145 (65.9) | - |  |
| Active disease ($\geq 5$) | 119 (36.7) | 44 (42.3) | 75 (34.1) | - | |
| SCCAI | 257 (88.9) | 84 (85.7) | 173 (90.6) | - | 0.14 |
| Remission ($\leq 2$) | 152 (59.1) | 44 (52.4) | 108 (62.4) | - | |
| Active disease (>2) | 105 (40.9) | 40 (47.6) | 65 (37.6) | - |  |
| **Surgical history** |  |  |  |  |  |
| Ileocecal resection, *n* (%) | 132 (20.6) | 40 (18.9) | 92 (21.5) | - | 0.47 |
| Colon resection (or partial), *n* (%) | 146 (22.8) | 55 (25.9) | 91 (21.3) | - | 0.19 |
| Small intestinal (partial) resection, *n* (%) | 82 (12.8) | 29 (13.7) | 53 (12.4) | - | 0.71 |

Data are presented as proportions *n* with corresponding percentages (%), mean ± standard deviation (SD) or as median [interquartile range, IQR] in case of continuous variables. *P*-values $\leq 0.05$ were considered statistically significant. <sup>†</sup>Use of TNF-α-antagonists included use of infliximab, adalimumab, golimumab and certolizumab pegol. Abbreviations: BMI, body-mass index; CD, Crohn's disease; HBI, Harvey-Bradshaw Index; IBD, inflammatory bowel disease; TNF-α , tumor necrosis factor alpha; SCCAI, Simple Clinical Colitis Activity Index; UC, ulcerative colitis.

### Mucosal gene expression reflects tissue specificity, inflammatory status and disease subtypes

Principal component analysis (PCA) showed that gene transcriptional patterns could be stratified by biopsy location (ileum vs. colon), inflammatory status (non-inflamed vs. inflamed) and IBD subtype (CD vs. UC) in the first two components (**Fig. 2A**), consistent with previous observations [13]. Tissue location and inflammatory status were significantly associated with the first two PCs (biopsy location, ileum vs. colon: *P*_Wilcoxon_=2.87x10^-12^; biopsy inflammatory status, *P*=7.15x10^-27^), whereas disease/control status (CD vs. UC vs. controls) was associated with the second PC (*P*=2.14x10^-16^).

**Figure 2.**
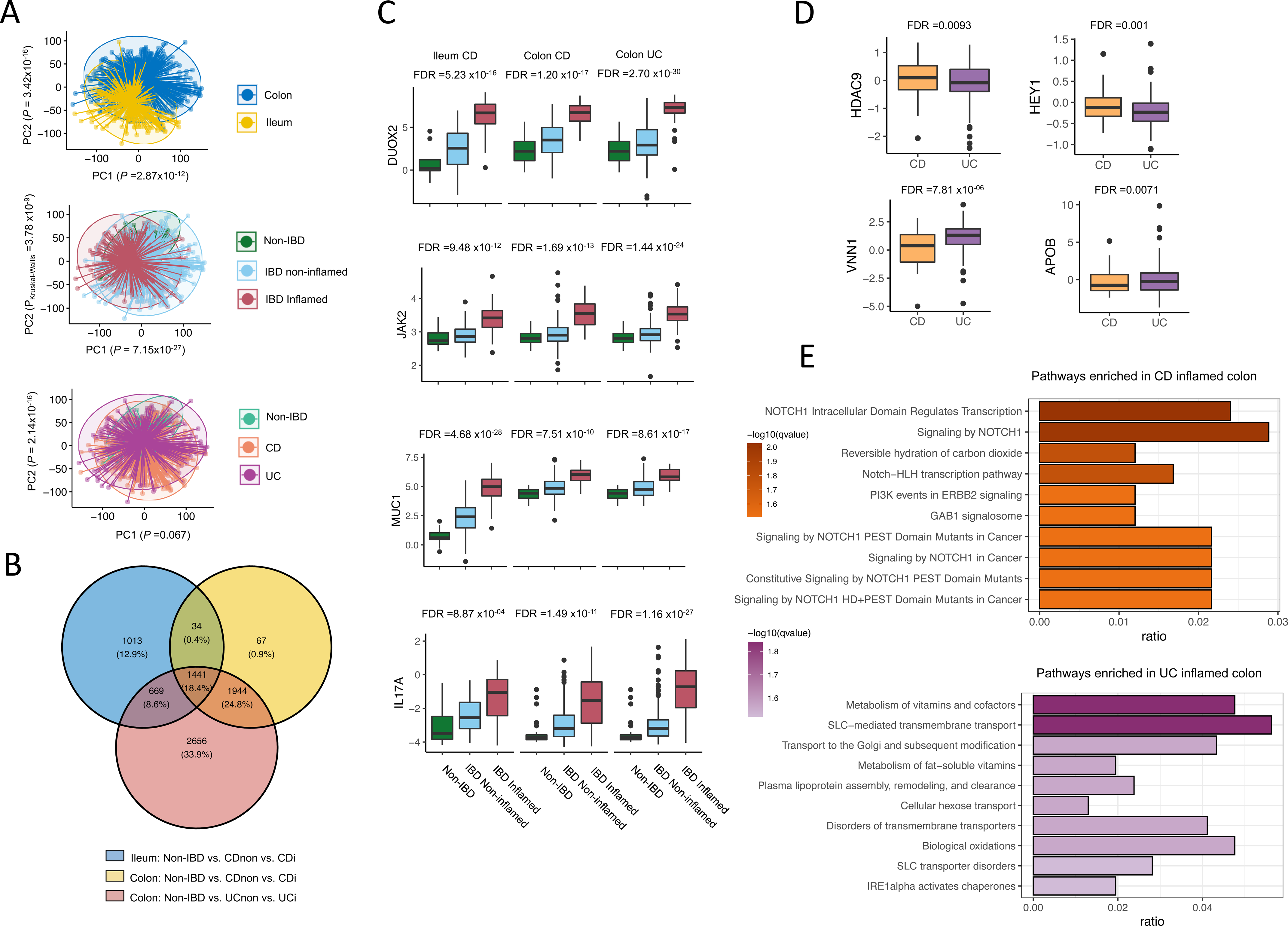
**Mucosal host gene expression patterns in intestinal tissue from patients with IBD and controls. a**, Principal component analysis, labeled by tissue location (ileum/colon), inflammatory status (non-inflamed/inflamed) and disease diagnosis (control/CD/UC), shows that variation in host gene expression can be significantly explained by tissue location and inflammatory status. **b**, Venn diagram of inflammation-associated genes from three comparisons: 1) ileal tissue from controls vs. non-inflamed tissue from patients with CD vs. inflamed tissue from patients with CD, 2) colonic tissue from controls vs. non-inflamed tissue from patients with CD vs. inflamed tissue from patients with CD and 3) colonic tissue from controls vs. non-inflamed tissue from patients with UC vs. inflamed tissue from patients with UC (all FDR <0.05). **c**, Relevant examples of four inflammation-associated genes, *DUOX2*, *JAK2*, *MUC1* and *IL17A*, illustrating the presence of tissue inflammation (FDR <0.05). **d**, Relevant examples of inflammation-associated genes differentially expressed between patients with CD and UC (keeping tissue location and inflammatory status constant) showing higher expression of *HDAC9* (histone deacetylase 9) and *HEY1* (hairy/enhancer-of-split related with YRPW motif protein 1) in patients with CD and higher expression of *VNN1* (pantetheinase) and *APOB* (apolipoprotein B) in patients with UC. **e**, Analysis of pathways associated with either the presence of CD (orange) or UC (purple) demonstrates that genes upregulated in CD are mainly associated with Notch-1 signaling, whereas pathways upregulated in UC are mainly related to vitamin and cofactor metabolism, SLC-mediated transmembrane transport and intracellular protein modification. Pathways were annotated using the Reactome pathway database. CDi, inflamed tissue from patients with Crohn’s disease. CD-non, non-inflamed tissue from patients with Crohn’s disease. FDR, false discovery rate. PC, principal component. UCi, inflamed tissue from patients with ulcerative colitis. UC-non, non-inflamed tissue from patients with ulcerative colitis.

Inflammation-associated gene expression showed overlap between inflamed biopsies from ileal CD, colonic CD and UC (**Fig. 2B**). Differential expression analyses between non-IBD controls, non-inflamed and inflamed biopsies in all these three groups revealed 3157, 3486, and 6710 differentially expressed genes (DEGs), respectively (FDR<0.05) (**Supplementary Table S1**). These DEGs fall mainly within interleukin signaling, neutrophil degranulation and extracellular matrix (ECM) organization pathways (FDR_Fisher_<0.05, **Extended Data Fig. S1**). Overlapping results from all three differential expression analyses identified 1437 shared DEGs, including *DUOX2*, *MUC1*, *JAK2*, *OSM* and *IL17A* (**Fig. 2C**). We also observed an enrichment of these DEGs in IBD- associated genomic loci (*P*_Fisher_=9.6x10^-9^) [2].

We then investigated the genes differentially expressed between inflamed colonic tissue from patients with CD and UC. In total, 1466 genes were differentially abundant, of which 733 (50%) were overrepresented in CD and 733 (50%) in UC (FDR<0.05) (**Supplementary Table S2**). Pathway enrichment analysis showed the Notch-1 signaling pathway (e.g. *HDAC9* and *HEY1*, **Fig. 2D**) to be highly upregulated in CD compared to UC, whereas vitamin, cofactor and lipoprotein metabolism pathways (e.g. *VNN1* and *APOB*, **Fig. 2D**) were more pronounced in UC (**Fig. 2E**), which corroborates previous findings [17–20]. Cell type–deconvolution revealed that plasma cells, endothelial cells and Th2-lymphocytes were significantly increased in UC compared with CD (FDR<0.05, **Supplementary Table S3**), suggesting that distinct immunological mechanisms are involved in CD and UC.

### Mucosal microbiota composition is highly personalized

The most common bacterial phylum observed across all tissue samples was Bacteroidetes (CD: 58%, UC: 58%, controls: 66%), followed by Firmicutes (CD: 27%, UC: 33%, controls: 23%) and Proteobacteria (CD: 14%, UC: 8%, controls: 9%). Interestingly, the overall mucosa-attached microbial composition was similar between colonic and ileal biopsies and independent of inflammation (**Extended Data Fig. S2**). Only seven bacterial taxa were differentially abundant between patients and controls (**Supplementary Tables S4-5**), consistent with previous findings [13,21,22].

Shannon diversity was significantly lower in samples from patients with CD compared to UC and non-IBD controls (*P*=2.75x10^-16^ and *P*=0.03, respectively, **Fig. 3A**). This difference was still present when comparing only colonic biopsies from patients with CD to those from UC, indicating that this difference was not solely attributable to ileal CD (**Extended Data Fig. S3**). Differences in microbial communities between tissue samples were evaluated by quantifying the Aitchison’s distance (**Fig. 3B-C**). We obtained comparable findings when we externally validated our results using data derived from the HMP2 cohort (**Extended Data Fig. S4**) [13].

**Figure 3.**
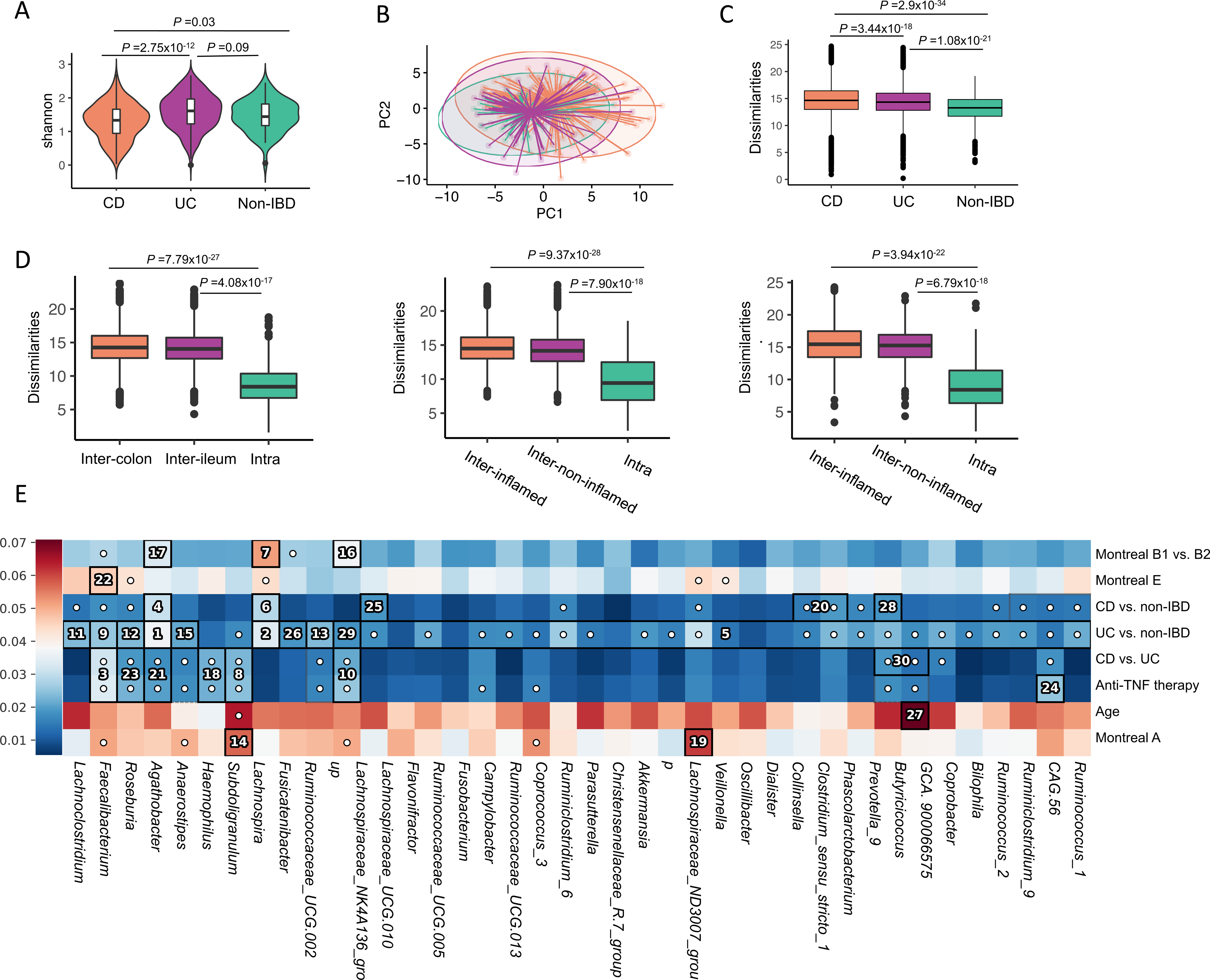
**Overall characterization of mucosa-attached microbiota in patients with IBD and controls. a**, Microbial alpha-diversity (Shannon index) was lowest in patients with CD (n=351) compared to patients with UC (n=289) and non-IBD controls (n=56). **b**, PCA plot based on Aitchison’s distances demonstrates the microbial dissimilarity of the mucosa-attached microbiota (colors as in **a**). **c**, Microbial dissimilarity (Aitchison’s distances) comparison between non-IBD control, CD and UC. Microbial dissimilarity is highest in biopsies from patients with CD, followed by patients with UC and non-IBD controls. **d**, Microbial dissimilarity is higher in samples from different individuals when compared to paired samples from the same individual, which includes paired inflamed–non-inflamed tissue from ileum and colon (left panel, inter-colon: *n*=11,430, inter-ileum: *n*=7,377, intra: *n*=203), paired colonic tissue samples from inflamed and non-inflamed areas (middle panel, inter-inflamed: *n*=7,372, inter-non-inflamed: *n*=8,369, intra: *n*=166) and paired ileal tissue samples from inflamed and non-inflamed areas (right panel, inter- inflamed: *n*=1,590, inter-non-inflamed: *n*=1,592, intra: *n*=73). **e**, Hierarchical analysis performed using an end-to-end statistical algorithm (HAllA) indicates the main phenotypic factors that correlate with intestinal mucosal microbiota composition. Heatmap color palette indicates normalized mutual information. Numbers or dots in cells identify significant pairs of features (phenotypic factors vs. bacterial taxa) in patients with IBD and controls. CD, Crohn’s disease. PCA, principal coordinate analysis. UC, ulcerative colitis.

**Figure 4.**
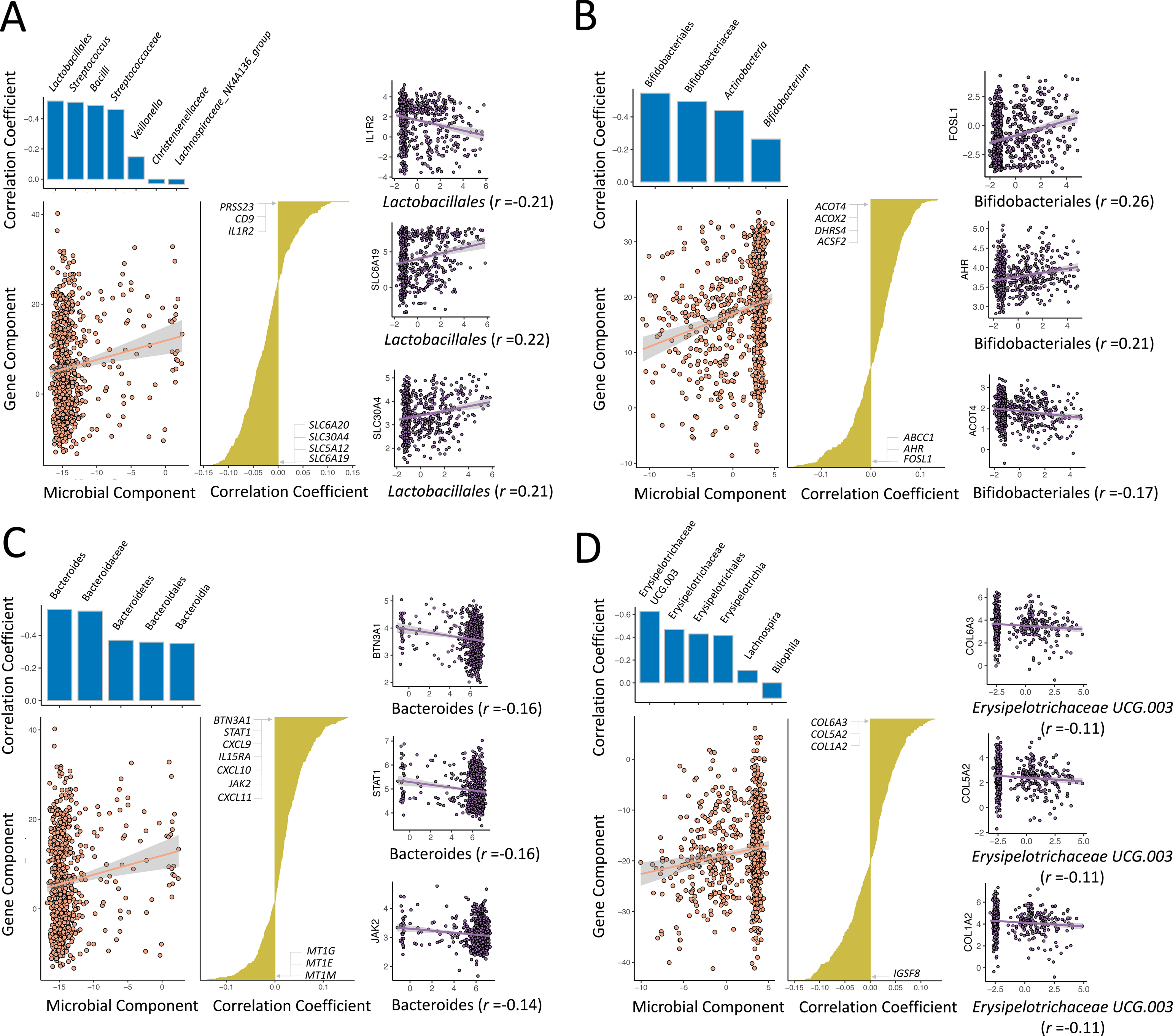
Mucosal host–microbe interaction modules in the context of IBD. Sparse canonical correlation analysis (sparse-CCA) was performed to identify distinct correlation modules of mucosal gene expression vs. mucosal microbiota through the identification of sparse linear combinations of two separate distance matrices that are highly correlated. Using 1437 inflammation-related genes and 131 microbial taxa as input, we identified six distinct pairs of significantly correlated gene–microbe components (FDR<0.05). **a**, A diverse group of mainly lactic acid producing bacteria (LAB) represented by order Lactobacillales, genus *Streptococcus*, class Bacilli, family *Streptococcaceae* and, to a lesser extent, genus *Veillonella*, family *Christensenallaceae* and the *Lachnospiraceae* NK4A136 group is associated with host pathways predominantly related to solute transport and liver metabolism. **b**, The abundance of mucosal *Bifidobacterium* bacteria is inversely associated with host fatty acid metabolism pathways (e.g. *ACOT4*, *ACOX2* and *ACSF2*) and positively associated with expression of specific genes, including *FOSL1*, *AHR* and *ABCC1/MRP1*. **c**, Mucosal *Bacteroides* bacteria inversely correlate with expression of genes representing host interleukin signaling pathways (e.g. *STAT3*, *JAK2*, *CXCL9* and *IL15RA*) but positively correlate with expression of genes representing metal ion response and metallothionein pathways (e.g. the metal ion response transcription factors *MT1G*, *MT1E* and *MT1M*). **d**, Mucosal *Erysipelotrichaceae* abundance inversely associates with expression of genes involved in collagen biosynthesis and collagen trimerization (e.g. *COL1A2*, *COL4A1* and *COL5A2*). Details of the two other significantly correlated pairs of components are presented in **Box 2**.

Intra-individual microbial dissimilarity was lowest in all our comparative analyses of paired tissue samples (**Fig. 3D**). Hierarchical clustering analysis performed on paired samples demonstrated a clear tendency of these samples to cluster together, a finding that we could also replicate in the HMP2 cohort data (**Extended Data Fig. S5A**) [13]. Overall, our data demonstrate that the composition of the mucosal microbiota is highly personalized and that inter-individual variability dominates over the effects of tissue location or inflammatory status.

We then aimed to identify phenotypic factors that shape the composition of the mucosal microbiota using Hierarchical All-against-All association (HAllA) analysis. This allowed us to study the relative associations between microbial taxa and phenotypic factors and disease characteristics (**Fig. 3E**, **Supplementary Table S6**). Analysis at bacterial genus level revealed that the main factors correlating with mucosal microbiota composition are stricturing disease in CD (fibrostenotic CD, Montreal B2), usage of TNF-α-antagonists, age at time of sampling, age of onset and the comparisons of patients with CD vs. controls, UC vs. controls and CD vs. UC. In contrast, inflammatory status and tissue location did not show a significant effect, and this was also the case within the HMP2 cohort data (**Extended Data Fig. S5B**). These findings are in line with several previous observations from which age at diagnosis, age at sampling and TNF-α-antagonist use emerged as critical determinants of mucosal microbiota composition [22].

### Distinct host–microbe interaction modules are identified in relation to IBD

To capture the main microbial taxa associated with inflammation-associated gene expression, we combined the data and performed sparse canonical correlation analysis (sparse-CCA) on 1437 inflammation-associated genes and 131 microbial taxa. This approach enabled us to identify gene pathways and groups of microbiota and their potential correlations. In total, we found six distinct pairings of groups of genes with bacterial taxa to be significantly correlated with each other (FDR<0.05, **Supplementary Tables S7-S18**). To prioritize the individual genes and bacteria involved in the sparse-CCA analysis, we performed individual pairwise gene–bacteria associations, which revealed 312 significant gene–bacteria pairs, with most pairs (94.17%) overlapping with the sparse-CCA results. We then replicated these associations in the HMP2 cohort (Spearman correlation ρ=0.16, *P*=0.005, **Supplementary Table S19, Extended Data Fig. S6, Methods**). Further details on the most intriguing individual pairwise gene– bacteria associations are discussed in **Box 1**.

### Mucosal lactic acid-producing bacteria positively correlate with nutrient uptake and solute transport

In the first significant pair of correlated components (component pair 1, *P*=5.72x10^-14^, FDR<0.05), the bacterial component is represented by bacteria from order Lactobacillales, family *Streptococcaceae*, class Bacilli and genus *Streptococcus* and, to a lesser extent, genus *Veillonella*, family *Christensenallaceae* and the *Lachnospiraceae* NK4A136 group (**Supplementary Tables S7-S8**). This bacterial component is mainly represented by lactic acid producing bacteria (LABs, including Lactobacillales, Bacilli, *Streptococcaceae*, *Streptococcus*) that actively participate in physiological food digestion, particularly carbohydrate fermentation, with lactic acid being their main metabolic product. Many of these bacterial groups are associated with genes involved in pathways related to solute transport and liver metabolism, including SLC-mediated transmembrane transport of bile salts, organic acids, metal ions and amine compounds; amino acid transport; biological oxidation; cytochrome P450 enzymes and the ephrin signaling pathway (involved in the migration of intestinal epithelial cells along the crypt-villus axis).

LABs are widely present in commercially available probiotics, and their beneficial effects on intestinal epithelial health are well-recognized [23]. SLC transporters mediate the bidirectional passage of nutrients such as sugars, amino acids, vitamins, electrolytes and drugs across the intestinal epithelium [24]. Although SLC transporters are often found to be dysregulated in patients with IBD (particularly CD), their expression may be stimulated and subsequently restored by commensal probiotic bacteria [25–27]. Taken together, however, we foresee that this host–microbe interaction component might not be IBD-specific as the genes and bacteria involved have important physiological functions in nutrient digestion and absorption.

### Mucosa-residing Bifidobacterium species show significant interplay with host fatty acid metabolism and bile acid transport pathways

The second pair of significantly associated components (component pair 3, *P*=1.89x10^-8^, FDR<0.05) is predominantly represented by bifidobacteria (**Supplementary Tables S9-S10**). The top associated pathways are represented by genes involved in fatty acid metabolism, including fatty acid biosynthesis (e.g. *ACOT4* and *ACSF2*), arachidonic acid metabolism (e.g. *CYP2J2* and *EPHX2*) and genes involved in peroxisomal protein import and fatty acid synthesis (e.g. *PEX5* and *ACOT4*), and these genes are all inversely associated with the bacterial component. In contrast, the genes *AHR* (encoding for the aryl hydrocarbon receptor) and *ABCC1* (encoding multidrug resistance protein 1) are positively correlated with the bacterial component. The inverse associations between bifidobacteria and the expression of genes involved in adipogenesis are consistent with findings from animal and small-scale human studies that investigated the effects of treatment with *Bifidobacterium* species on fatty acid metabolism [28–32]. Our findings may reflect the anti-inflammatory and anti-lipogenic role of bifidobacteria, which has previously been demonstrated in experimental settings, and may support the therapeutic potential of microbiome-directed interventions in attenuating or preventing colitis [31, 32].

### Mucosal Bacteroides associate with host interleukin signaling and metal ion response pathways

The third pair of significantly correlated components (component pair 7, *P*=1.28x10^-4^, FDR<0.05) is represented by Bacteroidetes. Twenty-four different pathways were significantly associated with this microbial component (**Supplementary Tables S11- S12**). A number of interferon signaling pathways (e.g. IFN-α, IFN-β and IFN-γ as well as the IL-2, IL-4, IL-6, IL-10, IL-12 and IL-13 signaling pathways) are all inversely associated with the microbial component. In addition, metal ion response and metallothionein pathways (e.g. metal ion transcription factors *MT1A*, *MT1E*, *MT1F*, *MT1G* and others) are positively associated with the microbial component. Taken together, these observations could suggest a predominance of potentially beneficial *Bacteroides* species associated with this component. Previous studies have shown that *Bacteroides* can exert either beneficial, mutualistic, or pathogenic effects on the host, depending on local interactions, intestinal location and nutrient availability [21, 33]. The co-occurrence of *Bacteroides* with lower expression of interleukin signaling pathways is supported by experimental work that found potential anti-inflammatory and protective roles for these bacteria in the context of intestinal inflammation [34,39,40]. Still, the relative contributions of each of these species, as well as their behavior in the context of intestinal inflammation, remains elusive, although our data might reflect an overrepresentation of anti-inflammatory members [34-39,43]. The positive associations between *Bacteroides* and expression of metal ion response genes and metallothioneins (MTs) are intriguing in the context of IBD because aberrant MT homeostasis and intracellular zinc metabolism have been implicated in disease pathophysiology [44–48].

### Mucosal Erysipelotrichaceae bacteria interact with collagen biosynthesis pathways

In the fourth pair of significantly correlated components (component pair 8, *P*=1.22x10^-4^, FDR<0.05), the microbial component, represented by the family *Erysipelotrichaceae*, is inversely associated with the expression of genes belonging to a wide range of ECM and collagen genes that are involved in collagen biosynthesis, integrin cell surface interactions, collagen chain trimerization, collagen fibril cross-linking, collagen fibril assembly, ECM proteoglycans, collagen degradation and related pathways (**Supplementary Tables S13-S14**). Similar to *Bacteroides*, the precise role of *Erysipelotrichaeae* in the context of IBD has not yet been fully elucidated. Some studies found lower abundances of *Erysipelotrichaceae* in patients with new-onset CD [49] and postoperative active CD [50], whereas others reported higher levels of *Erysipelotrichaceae* in the context of ileitis [51] and TNF-regulated CD-like transmural inflammation [52]. These inconsistencies have been suggested to be due to *Erysipelotrichaceae* behaving differently in response to intestinal inflammation, but they may also reflect incomplete characterization of the precise species that belong to the family of *Erysipelotrichaceae* [53]. Interactions between *Erysipelotrichaceae* and ECM/collagen remodeling pathways have not yet been reported in the context of IBD, but they would be particularly relevant because fibrosis occurs in a large fraction of patients with CD and *Erysipelotrichaceae* bacteria have been associated with fibrotic conditions beyond IBD [54–60].

### Patients with fibrostenotic CD exhibit a *Lachnoclostridium*-associated gene network involved in immune regulation

In pairwise comparative analyses, patients with fibrostenotic CD (Montreal B2, *n*=107) and patients using TNF-α-antagonists (*n*=113) exhibited several differentially abundant microbial taxa. We therefore analyzed microbiota-associated host mucosal gene interactions in these phenotypes (**Figure 5**). Pairwise comparisons between patients with non-stricturing, non-penetrating disease vs. fibrostenotic CD revealed 2639 differentially abundant genes that were enriched in cellular energy metabolism and immune system pathways (FDR<0.05, **Supplementary Table S20**). When comparing microbial taxa, abundances of mucosal *Faecalibacterium*, *Erysipelotrichaceae_UCG- 003* and *Coprococcus_3* were lower in fibrostenotic CD, whereas abundances of *Lachnoclostridium* and *Flavonifractor* were elevated in these patients (FDR<0.05). We hypothesized that these altered bacterial abundances and gene expression patterns may also translate into altered microbiota–gene networks relating to fibrostenotic CD. In patients with non-stricturing, non-penetrating CD, we observed 1508 individual gene– bacteria associations (corresponding to 84 different pathway–bacteria associations), whereas we found 541 individual associations (corresponding to 40 different pathway– bacteria associations) in patients with fibrostenotic CD. Comparing each bacteria-associated gene cluster between patients with non-stricturing, non-penetrating and fibrostenotic CD (FDR <0.05, **Methods, Supplementary Table S21**) identified four distinct networks represented by mucosal *Lachnoclostridium*, *Coprococcus*, *Erysipelotrichaceae* and *Flavonifractor*. The most significantly altered connections were associated with *Lachnoclostridium*, which was associated with 955 genes in patients with non-stricturing, non-penetrating CD, and these connections were mainly involved in cell activation pathways such as vesicle-mediated cellular transport and membrane trafficking (**Fig. 5A**). In total, 148 genes were associated with *Lachnoclostridium* in patients with fibrostenotic CD (FDR<0.05), and these genes were involved in cellular immunoregulatory interactions and adaptive immune system pathways (e.g. *CD8A*, *CLEC2B* and *CXCR5*), tyrosine kinase signaling (e.g. *FGF16*), opioid signaling and G alpha (s) signaling events (mediated via cAMP-dependent protein kinases, e.g. *POMC*, *GNG7* and *GNG11*) and vesicle-mediated transport (e.g. *APOE*, *COLEC12* and *KIF3B*) (**Fig. 5A-B**).

**Figure 5.**
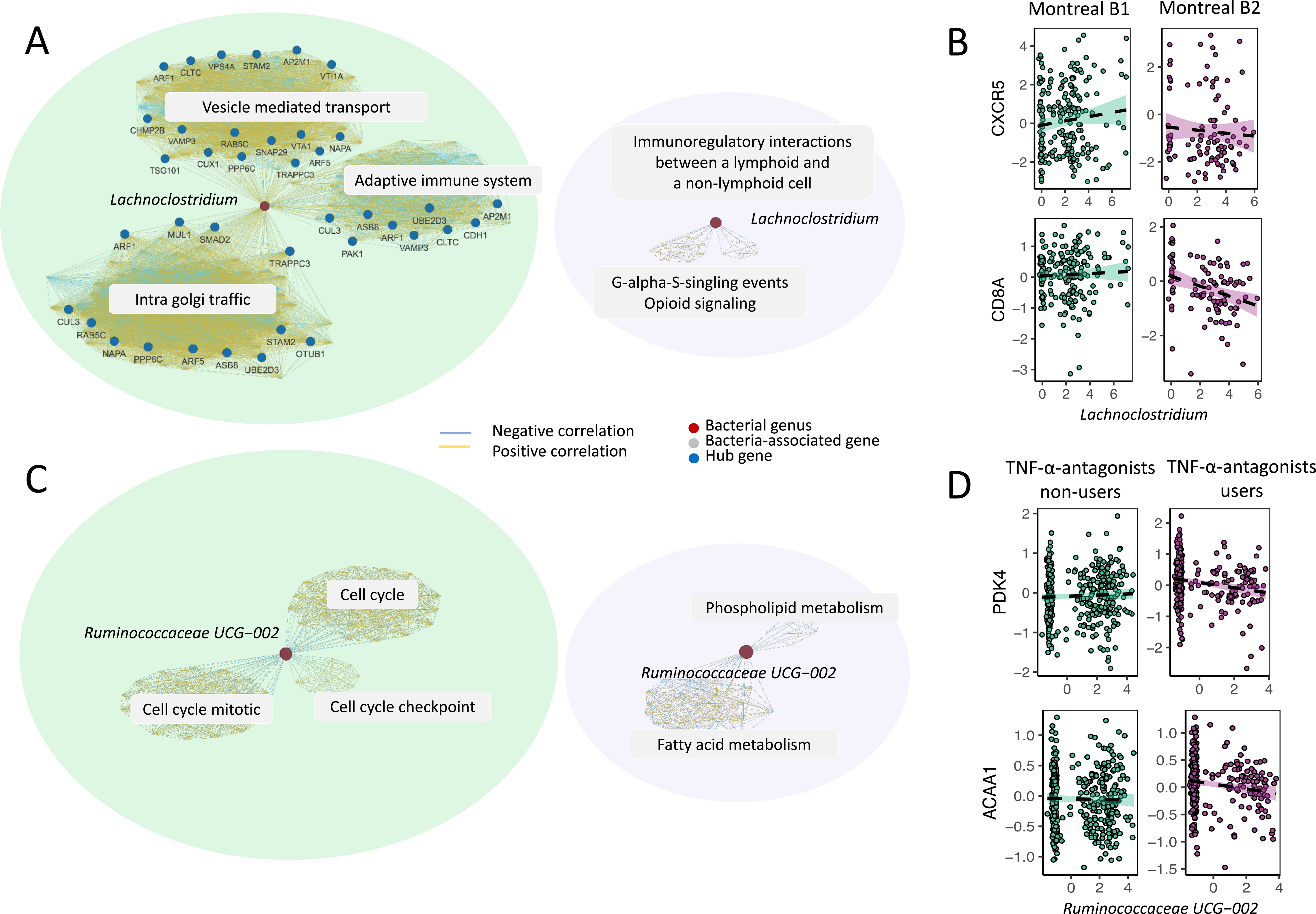
**Fibrostenotic CD and TNF-**α**-antagonist usage significantly alter mucosal host–microbe interactions in the context of IBD.** CentrLCC-network analyses were performed to characterize altered mucosal host–microbe interactions between different patient phenotypes. Overall, fibrostenotic CD (Montreal B2 vs. non-stricturing, non-penetrating CD, i.e. Montreal B1) and use of TNF-α-antagonists (vs. non-users) demonstrated significant modulation of observed mucosal host–microbe associations. **a**, Network graphs showing microbiota–gene association networks in patients with non-stricturing, non-penetrating CD (Montreal B1) (left) and patients with fibrostenotic CD (Montreal B2) (right). When comparing these patient groups, 5 bacterial taxa and 2639 host genes were significantly different (FDR<0.05). Four of the five bacterial taxa were significantly altered in fibrostenotic CD vs. non-stricturing, non-penetrating CD, and *Lachnoclostridium* was the top bacteria involved (covering 63% of total associations in non-stricturing, non-penetrating CD and decreasing to 27% in fibrostenotic CD). In general, patients with fibrostenotic CD were characterized by a loss of *Lachnoclostridium–*gene interactions. Red dots indicate gut microbiota. Blue dots indicate hub genes. Gray fields indicate the main pathways represented by the associated genes. Yellow lines indicate positive associations between gene expression and bacterial abundances. Light blue lines indicate negative associations. **b**, Key examples of *Lachnoclostridium–*gene interactions that were significantly altered in patients with fibrostenotic CD compared to patients with non-stricturing, non-penetrating CD, including genes involved in immunoregulatory interactions between lymphoid and non-lymphoid cells and tyrosine kinase signaling (*CD8A* and *CXCR5*). **c**, Network graphs showing microbiota–gene interaction networks in patients not using TNF-α-antagonists (left) vs. patients using TNF-α-antagonists (right). When comparing both groups, 3 bacterial groups and 513 genes were differentially abundant (FDR<0.05). Among these, a single bacterial group, represented by *Ruminococcaceae*_UCG_002, was altered in interactions with host genes in patients using TNF-α-antagonists. **d**, Key examples of *Ruminococcaceae–*UCG_002–gene interactions significantly altered in TNF-α-antagonists users vs. non-users. These genes were involved in general biological processes such as the cell cycle but also included genes involved in fatty acid metabolism (*PDK4* and *ACAA1*).

Earlier studies had shown that *Lachnoclostridium* bacteria are generally increased in patients with (complicated) CD, e.g. postoperative CD [61], ASCA-positive CD [62] and active granulomatous colitis [63]. Recently, *Lachnoclostridium* was also associated with non-invasive diagnosis of colorectal adenoma and colorectal cancer [64, 65]. These associations may potentially explain associations with genes involved in cellular proliferation and activation pathways. Increased abundances of *Lachnoclostridium* have been observed in relation to pulmonary fibrosis and its progression [66] but not in relation to intestinal fibrosis. In contrast, reduced abundances of *Faecalibacterium* and *Eubacterium* species (belonging to the *Erysipelotrichaceae* family) have previously been associated with luminal narrowing in patients with pediatric ileal CD [67]. Our results suggest that it is not only increased *Lachnoclostridium* abundances that may play a role in fibrostenotic CD, host immune-regulatory expression patterns may also vary along with these bacterial shifts. Notably, as the tissues investigated in our study were not derived from fibrotic regions, our findings show that these gene expression signatures are already present in non-stenotic intestinal tissue.

### Use of TNF-**α**-antagonists is associated with *Ruminococcaceae*-associated gene interactions related to fatty acid metabolism

Subsequently, we investigated the impact of TNF-α-antagonist use on mucosal host– microbe interactions. Pairwise comparisons revealed that TNF-α-antagonist use was significantly associated with three different bacterial taxa, *Faecalibacterium*, *Ruminococcaceae_UCG-002*, and *Ruminococcaceae_UCG-005* (all showing reduced abundances in users), and 513 different genes (FDR<0.05, **Supplementary Table S22**). For instance, one of the top genes associated with TNF-α-antagonist use was *CXCL13*, which encodes B cell attracting chemokine 1. By comparing each taxa-associated gene cluster between patients using and not using TNF-α-antagonists, we identified a single cluster represented by mucosal *Ruminococcaceae_UCG-002* that was significantly altered in users vs. non-users (FDR<0.05, **Supplementary Table S23**). *Ruminococcaceae_UCG-002* bacteria were associated with 135 genes in non-users, and these genes were mainly enriched in cell cycle–associated pathways (e.g. *PRIM1* and *PRIM2*), including mitosis-, prometaphase- and DNA-replication-associated genes (**Fig. 5C-D**). However, the *Ruminococcaceae_UCG-002*-associated genes in TNF-α-antagonist users (FDR<0.05) were predominantly involved in lipid/fatty acid metabolism (e.g. *ACAA1*, *ACSL5* and *PDK4*), glycerophospholipid biosynthesis and phospholipid metabolism. *Ruminococcaceae* comprise multiple distinct bacterial genera. Some of these are part of the healthy gut microbiome [68], but others are potentially pathogenic and commonly enriched in IBD [13, 69]. The *Ruminococcaceae UCG_002* group is classified under the *Oscillospiraceae* family, which consists of obligate anaerobic bacteria including *Faecalibacterium prausnitzii*. Depending on their micro- environment, *Ruminococcaceae* UCG_002 bacteria can produce short-chain fatty acids due to their fiber-metabolizing capacity [70–72]. The inverse associations between *Ruminococcaceae*_UCG_002 and genes involved in (peroxisomal) fatty acid oxidation in patients using TNF-α-antagonists might reflect a beneficial therapeutic modulation, i.e. a reduction of fatty acid oxidation and lipotoxicity, and possibly even attenuation of microbiota-induced intestinal inflammation [73–85].

### Mucosal host–microbe interactions depend on individual dysbiotic status

As patients with IBD have microbial dysbiosis compared to healthy individuals, we hypothesized that the strength and/or direction of the individual gene–bacteria interactions may depend on the microbial community (eubiosis vs dysbiosis). We therefore performed PCA on the microbiota data and calculated dysbiosis scores for all patients and controls, as represented by the median Aitchison’s distances to non-IBD controls (**Fig. 6A**). Patients with IBD demonstrated higher dysbiosis scores compared to controls (CD vs. non-IBD: *P*=5.1x10^-8^, UC vs. non-IBD: *P*=0.0015), but there was no clear difference between patients with CD and UC (**Fig. 6B**). When comparing patients with IBD above and below the 90^th^ percentile of dysbiosis scores [13], 204 individual gene–bacteria interactions showed significant dependence on microbial dysbiosis (**Fig. 6C, Supplementary Table S24**) (FDR<0.05). We also performed permutation tests, which confirmed that the significant interactions were not observed by chance (**Methods**, FDR<0.05). In one example of these interactions, expression of the *PLAUR* gene encoding for the urokinase plasminogen activator surface receptor was positively associated with *Lachnospiraceae* abundance, but this shifted to an inverse association when only considering individuals with a high degree of mucosal dysbiosis (90–100%) (*P*=1.69x10^-6^). The Ly6/PLAUR domain containing protein 8 (Lypd8) also functions as an antimicrobial peptide and has previously been shown to be capable of protecting the host from invading pathogenic flagellated bacteria [86]. Another example is the positive association between *S100A8*, which encodes S100 calcium-binding protein A8 (also known as calgranulin A), and *Lachnospiraceae*, which showed a negative association in individuals with high dysbiosis (*P*=1.78x10^-5^). S100A8 has a wide variety of functions in regulating inflammatory processes and forms a heterodimer with S100A9, also known as calprotectin, which is used a as biomarker for inflammatory activity in IBD. Its known antimicrobial activity towards bacteria via chelation of zinc ions, which are essential for microbial growth, may therefore be disrupted in a dysbiotic environment [44]. Similar to the two previous examples, the observed association between the expression of *IL1RN* (encoding for the interleukin-1 receptor antagonist protein) and *Lachnospiraceae* shifted from positive to negative (*P*=4.10x10^-5^), indicating that the natural protection against the proinflammatory effects of IL-1β, which associates with *Lachnospiraceae* abundance, may be lost in circumstances of high microbial dysbiosis. Similarly, expression of the *CXCL17* gene encoding for a mucosal chemokine protein known to exert broad antimicrobial activity [87] positively correlated with *Lachnospiraceae* abundance, which was clearly different among individuals with higher dysbiosis scores.

**Figure 6.**
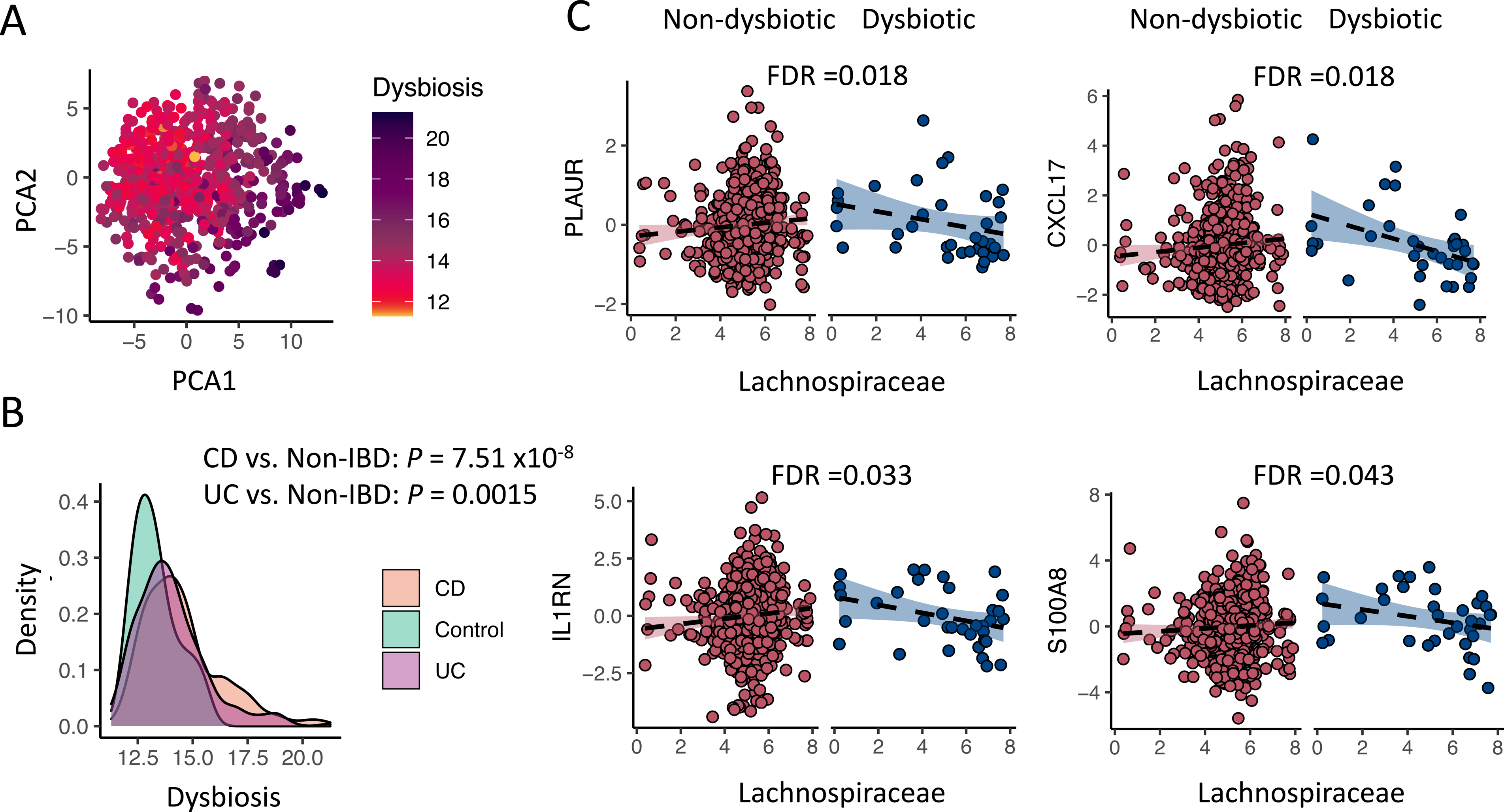
**Mucosal host–microbe interactions depend on individual dysbiotic status. a**, PCA of mucosal 16S rRNA sequencing data shows that degree of mucosal dysbiosis explains a large part of microbial variation. **b**, Dysbiosis scores were generally higher among patients with CD and UC compared to controls. **c**, Key examples of individual gene–bacteria interactions that demonstrate a directional shift upon higher dysbiosis (90–100%) as compared to patients with lower dysbiosis scores (0–90%). Mucosal *Lachnospiraceae* bacteria positively associate with the expression of the *PLAUR*, *CXCL17*, *IL1RN* and *S100A8* genes, whose gene products all have beneficial antimicrobial activity towards pathogenic bacteria. CD, Crohn’s disease. PCA, principal component analysis. UC, ulcerative colitis.

### Mucosal microbiota associate with variation in intestinal cell type–enrichment

Subsequently, we aimed to evaluate which intestinal cell types are involved in mucosal host–microbe interactions (**Figure 7, Extended Data Fig. S7**). Deconvolution of host gene expression data revealed that the mucosal microbiota was significantly associated with several cell types, but most evidently with intestinal epithelial cells, M1 macrophages, NK cells and mucosal eosinophils. Tissue inflammatory status and location also strongly contributed to the variation in most intestinal cell types. Mucosal microbiota that were significantly associated with intestinal epithelial cell enrichment typically belonged to the Firmicutes phylum, including *Agathobacter*, *Dialister*, *Lachnospira*, *Lachnoclostridium* and *Ruminococcaceae* (**Supplementary Table S25**).

**Figure 7.**
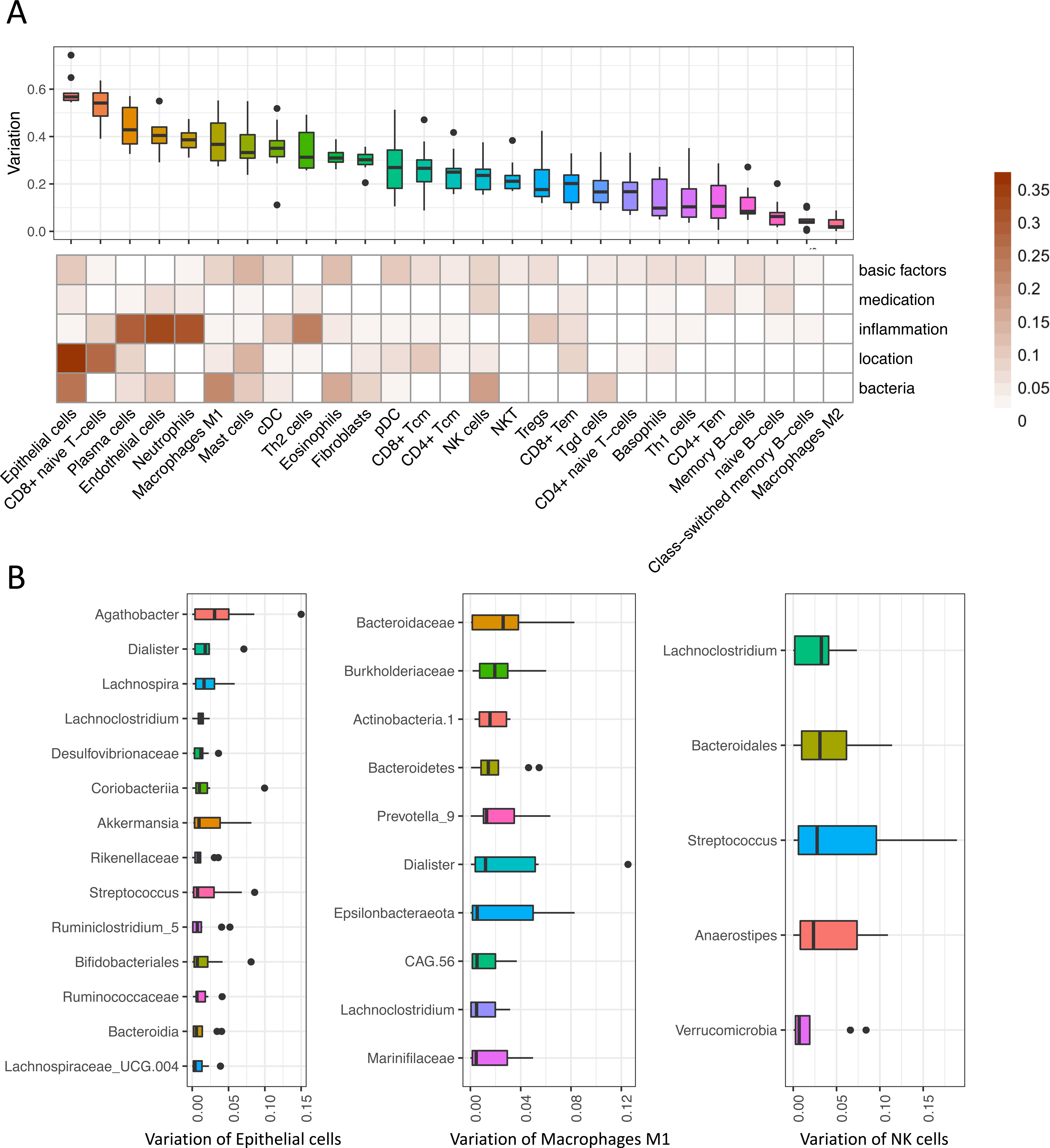
**Mucosal microbiota associate with distinct intestinal mucosal cell types**. **a**, Boxplots show the amount of variation in intestinal cell type–enrichment that could be explained by mucosal microbiota. Heatmap shows the contribution of other fitted models in explaining intestinal cell type–enrichment, including ‘basic factors’ (age, sex and BMI), medication use, tissue inflammatory status and tissue location. Mucosal microbiota contributed most to the variation in enrichment of intestinal epithelial cells, M1-macrophages, NK cells and eosinophils. **b**, Boxplots showing the contribution of the main bacterial taxa that explain the variation in mucosal enrichment of intestinal epithelial cells, M1-macrophages and NK cells—the cell types that interacted most strongly with the mucosal microbiota.

## Discussion

In this study, we show distinct mucosal host–microbe interactions in intestinal tissue from patients with IBD. Mucosal gene expression patterns in IBD are mainly determined by tissue location and inflammatory status and systematically demonstrate upregulation of distinct inflammation-associated genes, even in endoscopically non-inflamed tissue. Subsequently, we observed that the mucosal microbiota composition in patients is marked by high inter-individual variability. The main focus of our analyses, however, was integrative analysis of both data entities, which allowed us to comprehensively uncover many host–microbe associations, both on component level and as individual associations in IBD. Furthermore, we identify specific transcriptional networks that are significantly altered in patients with fibrostenotic CD and patients using TNF-α- antagonists and observe that these associations depend on the degree of mucosal dysbiosis. Finally, we show that mucosal microbiota are significantly associated with intestinal cell type composition, in particular with epithelial cells, macrophages and NK cells.

Tissue location and inflammatory status have the greatest impact on the variation in mucosal gene expression patterns. Enriched genes are mainly represented by those involved in pathophysiological pathways relevant to IBD, e.g. interleukin and interferon signaling and ECM remodeling. Patients with CD and UC show striking differences, e.g. Notch-1 signaling pathways are upregulated in CD, while genes involved in nutrient absorption and lipid metabolism are downregulated. Activation of Notch-1 signaling has been associated with improved mucosal barrier function, driven by lamina propria- residing CD4^+^-T-lymphocytes that induce intestinal epithelial cell differentiation [17].

Notch-1 signaling more efficiently spreads within CD intestinal epithelia, as compared to UC or control epithelia. Notch-1 is not only implicated in IBD, it also confers protection against the development of colorectal carcinoma via p53 signaling, thereby promoting cell cycle arrests and cellular apoptosis [18,88,89]. Since UC patients with long-lasting colonic inflammation have a higher risk of developing IBD-associated colorectal carcinoma, we hypothesize that downregulation of Notch-1 in these patients may potentially be involved in carcinogenesis.

Analysis of mucosal microbiota in patients with IBD reveals reduced alpha-diversity, microbial dissimilarity and marked intra-individual variability that is particularly strong in CD but still present to a lesser extent in UC. Given the large heterogeneity in IBD and the fact that compositional differences are largely attributable to individual phenotypic factors, cautious interpretation is warranted when associating mucosal microbial profiles to disease phenotypes or outcomes, rendering them inappropriate for diagnostic purposes. These observations corroborate those of previously published mucosal 16S studies in IBD [13,21,22]. Moreover, our findings align with a recent prospective meta-analysis study that concluded there is sparse evidence for additional population structure in mucosal microbiomes in IBD, e.g. microbiota-driven discrete disease subtypes within IBD [90].

Sparse-CCA analysis was performed to capture the key pathway–bacteria interactions. These include numerous inverse associations between bifidobacteria and expression of genes involved in fatty acid metabolism, which align well with previously published data from animal studies demonstrating anti-inflammatory and anti-lipogenic effects of *Bifidobacterium* treatment on chemically-induced intestinal inflammation [29,31,32]. For example, treatment with *Bifidobacterium adolescentis* IM38 attenuated high fat diet– induced colitis in mice by inhibiting lipopolysaccharide production, NF-κB activation and TNF-expression in colonic epithelial cells [31]. Likewise, treatment with *Bifidobacterium infantis*, with or without a combination of inulin-type fructans, ameliorated DSS-induced colitis in rats, as evidenced by decreased expression of IL-1β, malondialdehyde (MDA, a lipid peroxidation marker), decreased bacterial translocation and increased production of short-chain fatty acids [32]. In line with our findings, this supports the ongoing quest for efficacious probiotic (bifidobacteria-containing) interventions in patients with IBD [91, 92]. In addition, we observe a *Ruminococcaceae*-UCG-002-associated network of genes involved in (peroxisomal) fatty acid oxidation and lipotoxicity, which are inversely associated with these bacteria in patients using TNF-α-antagonists. Interestingly, multiple studies have observed that *Ruminococcaceae* increase after anti-TNF therapy in patients with CD and UC [73,75–77]. One of these studies specifically identified an association between the *Ruminococcaceae_*UCG-002 group and responsiveness to TNF-α-antagonists, albeit not in relation to host gene expression patterns [75].

Strikingly, many of the network-associated genes we observe are controlled by the PPAR-γ transcription factor, a butyrate sensor that may result in reduced lipotoxicity and reduced intestinal inflammation through prevention of overgrowth of potentially pathogenic bacteria [79–85]. These findings underscore the potential relevance of PPAR-γ as a therapeutic target in IBD [85].

We also observed an intriguing inverse relationship between *Erysipelotrichaceae* and intestinal ECM remodeling pathways, which may support the notion that intestinal fibrosis in IBD is highly linked to microbial composition [60,93,94]. Interestingly, a decreased relative abundance of *Erysipelotrichaceae* has previously been observed in patients with collagenous colitis [55] and cystic fibrosis–related lung fibrosis [56–58], as well as in mice with liver fibrosis and hepatocyte-specific *NOD2* deletions [59]. In CD, several bacterial species belonging to *Erysipelotrichaceae*, including *Clostridium innocuum* and *Erysipeloclostridium ramosum*, have been associated with the expansion of mesenteric adipose tissue (“creeping fat”), a unique feature of CD [60]. Creeping fat in CD has previously been characterized by higher abundances of *Erysipelotrichaceae* compared to adjacent mesenteric adipose tissue and underlying mucosal tissue and is accompanied by higher expression of ECM- and collagen-related genes. *C. innocuum* translocated to mesenteric fat, promoted fibrosis and stimulated tissue-remodeling in patients with CD, resulting in an adipose tissue barrier that may prevent systemic translocation of intestinal bacteria [60]. This phenomenon could potentially explain the inverse associations we observe between expression of ECM remodeling and mucosal *Erysipelotrichaceae*. In our differential network analyses, we observe a substantial decrease of *Lachnoclostridium*-associated genes in patients with fibrostenotic CD that are mainly associated with cellular immunoregulatory interactions and adaptive immune system pathways. These findings suggest that *Lachnoclostridium*-associated immunoregulatory expression patterns may play a role in fibrostenotic CD. Although little is known about the exact role of *Lachnoclostridium* in IBD, these bacteria were recently strongly associated with the development of colorectal cancer and with pulmonary fibrosis [64–66].

Another key host–microbe interaction module pertains to *Bacteroides*, which inversely correlates with interleukin signaling and positively associates with metal stress response transcription factors encoding for MTs. To maintain cellular redox balance, MTs detoxify heavy metal ions and scavenge ROS, thereby attenuating oxidative stress. Previous studies have shown that MTs may prevent experimental colitis or act as danger signals by mediating immune cell infiltration in the intestine [45, 46]. Although experimental evidence seems to be inconclusive, there is ample evidence indicating a role for aberrant MT homeostasis in IBD [47]. This mechanism depends on the intracellular accumulation of zinc, which induces autophagy under chronic NOD2-stimulation. In IBD, the mucosal microbiota may contribute to the regulation of MT expression, intracellular zinc homeostasis and autophagy, thereby regulating intracellular bacterial clearance by intestinal macrophages. Findings from this study may support a putative role for *Bacteroides* in modulating MT activation, thereby contributing to intracellular redox homeostasis, zinc levels, macrophage autophagy, or even host defense against pathogens. Importantly, MTs and zinc regulation constitute potential therapeutic targets in IBD [44-47, 95-97].

Individual gene–bacteria association analysis revealed distinct mucosal host–microbe interactions that largely overlap with those from the sparse-CCA analysis, but these provide more granular insight into the observed associations. Key examples of individual host gene–bacteria interactions are listed in **Box 1**. Amongst others, we demonstrate several host–microbe interactions that are putatively involved in immunological tolerance and prevention of autoimmunity (e.g. bifidobacteria and *FOSL1*/*KLF2* expression), colorectal carcinogenesis (e.g. *Anaerostipes* and *SMAD4*, *Akkermansia* and *YDJC*) and inflammatory signaling (e.g. *Oscillibacter* and *OSM* expression). Notably, many of these associations are dependent on fibrostenotic disease, TNF-α-antagonist use and the degree of mucosal dysbiosis. In addition, deconvolution of the mucosal RNA-seq data reveal cell type–specific patterns of microbial interactions that warrant further study, for example through single-cell RNA-seq studies.

Mucosal host–microbiota interactions have been investigated previously in both cohort (e.g. the HMP2 and Irish IBD) and experimental studies [12–16]. Alongside several observations consistent with previous findings, we identify many novel host–microbe interactions. Differences in sample size, patient phenotypes and sample handling may be at least partially responsible for these observations. In our study, large groups of gene–bacteria associations are revealed that cover a wide range of molecular mechanisms potentially relevant in the context of IBD, including immune response pathways, cellular processes and a variety of metabolic pathways. Moreover, our study features the largest sample size so far [12–15], and this enabled us to perform an integrative analysis with respect to the large disease heterogeneity and identify novel host–microbiota crosstalk related to different clinical characteristics. However, several limitations also warrant recognition. As our study is of cross-sectional origin, we cannot assess the longitudinal dynamics of host–microbe interactions to discover signatures for therapy responsiveness or disease prognosis. Consequently, our associative results cannot establish potential causality between microbial abundances and host gene expression. Functional experiments are thus required to validate the biological relevance of the individual host–microbe interactions, as well as their behavior in microbial ecosystems. Finally, bowel preparation prior to the endoscopic procedure or cross-contamination between biopsy sites during endoscopy can affect the mucosal microbiota composition [21,50,98].

Our results demonstrate a complex and heterogeneous interplay between mucosal microbiota and mucosal gene expression patterns that is concomitant with the strong impact of specific patient traits in a large cohort of patients with IBD. Our findings may guide development of mechanistic studies (e.g. host–microbe co-culture systems) that could provide functional confirmation of relevant pathophysiological gene–bacteria interactions and serve as a resource for rational selection of therapeutic targets in IBD. This study presents a large-scale, comprehensive landscape of intestinal host–microbe interactions in IBD that could aid in guiding drug development and provide a rationale for microbiota-targeted therapy as a strategy to control disease course. Future studies are warranted to focus on the integration of host–microbe interaction modules in prospective clinical trials investigating their utility for predicting disease course and responsiveness to treatment and for stratifying patients to facilitate therapeutic decision- making.

## Methods

### Study population

Patients with an established diagnosis of IBD were included at the outpatient clinic of the University Medical Center Groningen (UMCG) based on their participation in the 1000IBD project, for which detailed phenotypic information and multi-omics profiles had been generated [99]. Patients included in this study were at least 18 years old and were enrolled from 2003–2019. Diagnosis of IBD was based upon clinical, laboratory, endoscopic and histopathological criteria, with the latter criteria also used to determine the inflammatory status of collected biopsies. Detailed phenotypic data were collected for all patients, including age, sex, BMI (body weight divided by squared height), smoking status, Montreal disease classification, medication usage, history of surgery, clinical disease activity and histological disease activity, and all were assessed at time of sampling. Montreal disease classification was recorded from the closest visit to the outpatient clinic at time of sampling. Clinical disease activity was established using the Harvey-Bradshaw Index (HBI) for patients with CD and the Simple Clinical Colitis Activity Index (SCCAI) for patients with UC. We further included 17 healthy non-IBD controls (*n*=59 biopsies) who underwent endoscopy because of clinical suspicion of intestinal disease or within the context of colon cancer screening. All participants provided written informed consent prior to sample collection. This study was approved by the Institutional Review Board (IRB) of the UMCG, Groningen, the Netherlands (in Dutch: ‘Medisch Ethische Toetsingscommissie’, METc; IRB nos. 2008/338 and 2016/424) and was conducted according to the principles of the Declaration of Helsinki (2013).

### Mucosal RNA-sequencing

711 intestinal biopsies from 420 patients with IBD were collected. These were immediately snap-frozen in liquid nitrogen by an endoscopy nurse or research technician present during the endoscopic procedure. Biopsy inflammatory status was assessed based on histological examination by certified pathologists. Biopsies were stored at -80℃ until further processing.

RNA isolation was performed using the AllPrep DNA/RNA mini kit (Qiagen, reference number: 80204) according to manufacturer’s instructions. Homogenization of intestinal biopsies was performed in RLT lysis buffer including β-mercaptoethanol using the Qiagen Tissue Lyser with stainless steel beads (diameter 5 mm, reference number: 69989). For the first sample batch, sample preparation was executed using the BioScientific NEXTflex^TM^ Rapid Directional RNA-Seq Kit (Perkin-Elmer). Paired-end sequencing of RNA was performed using the Illumina NextSeq500 sequencer (Illumina). For the second sample batch, sample preparation was performed for construction of the Eukaryotic Transcriptome Library (Novogene). Paired-end sequencing of RNA was performed using the Illumina HiSeq PE250 platform. Sequencing was performed in two different batches, which necessitated pseudo-randomization (covering type of IBD diagnosis, biopsy location and disease activity) across plates to mitigate potential batch effects. The batch effects have been taken into account in all the analysis. On average, approximately 25 million reads were generated per sample.

Raw read quality was checked using FastQC with default parameters (ref v.0.11.7). Adaptors identified by FastQC were clipped using Cutadapt (ref v1.1) with default settings. Sickle (ref v1.200) was used to trim low-quality ends from the reads (length <25 nucleotides, quality <20). Reads were aligned to the human genome (human_glk_v37) using HISAT (ref v0.1.6) (with maximum allowance of two mismatches), and read sorting was performed using SAMtools (ref v0.1.19). SAMtools flagstat and Picard tools (ref v2.9.0) were used to obtain mapping statistics. Six samples with low percentage read alignment (< 90%) were removed. Gene expression was estimated using HTSeq (ref v0.9.1), based on Ensemble version 75 annotation, resulting in a RNA expression dataset featuring 15,934 genes. Expression data on gene level were normalized using a trimmed mean of *M* values, and *clr* transformation was applied, resulting in 826 mucosal RNA-seq samples.

### Mucosal 16S rRNA gene sequencing

Total DNA extraction of intestinal biopsies using 0.25 g of sample was performed as described previously, with minor modifications [100]. Microbial composition of intestinal biopsies was determined by Illumina MiSeq paired-end sequencing of the V3-V4 hypervariable region of the 16S rRNA gene (MiSeq Benchtop Sequencer, Illumina Inc., San Diego, USA). Amplification of bacterial DNA was performed by PCR using modified 341F and 806R primers with a six-nucleotide barcode on the 806R primer for multiplexing [101, 102]. Sequences of both primers can be found in **Supplementary Table S1**. Both primers contain an Illumina MiSeq adapter sequence, which is necessary for flow cell–binding in the MiSeq machine. A detailed overview of the PCR, DNA clean-up and MiSeq library preparation using a 2x300 cartridge can be found in the **Supplementary Methods**. Read trimming and filtering was done using *Trimmomatic* (0.33) to obtain an average read quality of 25 and a minimum length of 50. Quality was further checked using R package *DADA2* (v1.03) with the following parameters: truncLen=c(240,160), maxN=0, maxEE=c(2,2), truncQ=2 and rm.phix=TRUE. After error correction and chimera removal, the amplicon sequence variants were assigned to the *silva* database (v.132). Samples with >2,000 mapped reads were used for further analysis, resulting in 755 mucosal 16S samples. After accounting for overlap between mucosal RNA-seq and mucosal 16S data, 696 intestinal biopsies from 337 different patients and 16 non-IBD controls were available for host– microbiota interaction analyses.

### Statistical analysis

#### Descriptive statistics

Descriptive data are presented as means ± standard deviation (SD), medians [interquartile range, IQR] or proportions *n* with corresponding percentages (%). Between-group comparisons were performed using Mann-Whitney *U*-tests, Pearson’s chi-squared tests or Fisher’s exact tests (if *n* observations were <10). Nominal *P*-values ≤ 0.05 were considered statistically significant.

#### Mucosal gene expression analysis

Sample gene expression dissimilarity was calculated using Aitchison’s distances. General linear models were used to assess the associations between mucosal gene expression and clinical phenotypes while controlling for potential confounders, which were determined from our previous study (medication included the use of aminosalicylates, thiopurines and steroids) [103]. In particular, to assess the effect of mucosal inflammation on gene expression, we re-coded the inflammation status in an ordinal fashion as 0, 1 or 2 to represent biopsies from non-IBD controls, biopsies from non-inflamed tissue of patients with IBD and biopsies from inflamed areas of patients with IBD, respectively. Intestinal inflammatory status was thus treated as a continuous variable to account for presence of residual inflammation in biopsies marked as being taken from non-inflamed areas in the intestines. A correction for multiple hypotheses testing was applied using an FDR threshold of 5%.

(1) Inflammation-associated genes were identified in three comparisons: (1) CD colonic inflamed tissue vs. CD colonic non-inflamed tissue vs. non-IBD colonic tissue, (2) CD ileocecal inflamed tissue vs. CD ileocecal non-inflamed tissue vs. non-IBD ileocecal tissue and (3) UC colonic inflamed tissue vs. UC colonic non-inflamed tissue vs. non-IBD colonic tissue:

Gene ∼ intercept + inflammation + age + sex + BMI + medication + batch

(2) Clinical phenotype–associated genes were identified using the following model:

Gene ∼ intercept + Montreal/anti-TNF therapy + age + sex + BMI + inflammation + tissue location + medication + batch

#### Microbial characterization

Microbial richness and evenness was determined by calculating the Shannon index representing alpha-diversity of the gut microbiota. Microbial dissimilarity of samples was determined by calculating Aitchison’s distances after *clr* transformation using the R package *Compositions* (v2.02). Analysis of paired samples from the same individuals was performed while comparing microbial features between inflammation status,disease location and disease subtype using paired Wilcoxon tests. Factors potentially influencing mucosal microbiota were determined using Hierarchical All-against-All significance testing (HAllA) [104]. Associations between microbial features and biopsy inflammatory status, IBD diagnosis, disease location (biopsy origin) and clinical phenotypes were performed using general linear models (see below). Per sample, the mucosal dysbiosis score was defined as the median Aitchison distance from that sample to a reference sample set of non-IBD controls. Dysbiotic status was defined as being at the 90^th^ percentile of this score [13].

(1) Associations between microbial taxa and biopsy inflammation/location:

Taxa ∼ intercept + inflammation + location + age + sex + BMI + medication + batch + surgical resection

(1) Associations between microbial taxa and clinical phenotypes:

Taxa ∼ intercept + Montreal/anti-TNF therapy + inflammation + location + age + sex + BMI + medication + batch + surgical resection *Gene–microbiota interaction analysis*

We first focused on host inflammation-related genes (*n*=1,437) to investigate their potential associations with mucosal microbiota. Group-level correlations between gene expression and mucosal microbiota were performed using sparse-CCA using the residuals of genes and microbiota after correcting for age, gender, BMI, inflammation, tissue location and surgical resection separately. Sparse-CCA identifies the PCs from two related datasets that maximize the correlation between the two components. A set of enriched host pathways for all significant components was combined while adjusting for multiple comparisons using the FDR approach. Individual pairwise gene–microbiota associations were assessed by fitting a general linear model while adjusting for age, sex, BMI, inflammation status, tissue location, sequencing batch and medication use (including the use of aminosalicylates, thiopurines and steroids, see below). A gene– microbiota network analysis was visualized using the R package *ggview*.

(1) Individual gene–bacteria associations were determined using the following model:

Gene ∼ intercept + taxa + inflammation + location + age + sex + BMI + medication + batch

Second, we focused on host–microbiota interactions associated with fibrostenotic CD and usage of TNF-α-antagonists. Genes and taxa that were differentially abundant between clinical phenotypes were selected and then served as input for CentrLCC- network analysis using the *NetCoMi* R package. Hub nodes were defined as those with an eigenvector centrality value above the empirical 95% quantile of all eigenvector centralities in the network. This analysis was done in different groups separately (e.g. users and non-users of TNF-α-antagonists). To assess whether the taxa-associated gene networks were altered between groups, the associated genes for each taxa node were ranked within the total geneset background based on Z-scores. The Wilcoxon test was used to compare the two gene rank lists for each taxa.

Third, we assessed whether gene–microbiota associations depend on intestinal dysbiosis by modeling these associations using an additional interaction term in linear models. The dysbiosis score was treated as a continuous value. To determine whether these interactions were observed by chance, we also performed permutation tests that randomly shuffled the dysbiosis score 100 times across all samples, and then repeated the interaction models. On average, only three FDR-adjusted significant results were obtained for each round of permutation testing, suggesting that the rate of total false positives was approximately ∼ 0.014 (3/204).

2) *Gene ∼ intercept + taxa + dysbiosis + taxa * dysbiosis + inflammation + location + age + sex + BMI + medication + batch*

Fourth, enrichment of specific intestinal cell types was inferred from the RNA-seq data using the *Xcell* package in R. The effects of tissue location, inflammatory status and type of IBD diagnosis on expression levels of mucosal cell types were assessed using linear models, adjusting for age, sex, BMI, batch and medication usage. Subsequently, we used the *glmnet* R package to investigate the variation of cell type–enrichment that could be explained by the mucosal microbiota using *lasso* regression while employing a nested 10-fold cross-validation using six models:

1. Cell enrichment ∼ age + gender + BMI + batch
2. Cell enrichment ∼ medication (aminosalicylates, thiopurines, steroids, biologicals)
3. Cell enrichment ∼ inflammation
4. Cell enrichment ∼ tissue location
5. Cell enrichment ∼ bacteria abundance
6. Cell enrichment ∼ full factors mentioned above

The percentage of explained variance (R^2^) was calculated to estimate the variation in cell type–enrichment explained by the mucosal microbiota. All analyses were corrected for multiple testing using a FDR significance threshold of 0.05. All gene pathway enrichment analyses were conducted using the Reactome database from MsigDB [105, 106].

#### Replication in the HMP2 dataset

RNA-seq and 16S raw data were obtained from https://ibdmdb.org and reprocessed using the same pipeline in this study. After harmonizing with the phenotype file, we included 152 intestinal biopsies from the 85 patients with CD, 46 patients with UC and 45 non-IBD controls. First, gene expression and mucosal microbiota patterns were compared separately between this study and HMP2. Second, given the limited overlap in clinical phenotypes between the two cohorts, we restricted the replication analysis to inflammation-related host–microbiota interactions. Individual gene–microbiota associations were calculated using the same linear models used in this study while adjusting for age, gender, tissue location and inflammation status. Spearman correlation coefficients were used to assess the concordance between the Z-scores of gene– microbiota associations from the two studies.

## Acknowledgements

All authors would like to express their gratitude towards all participants of the 1000IBD cohort. In addition, the authors would like to thank Kate McIntyre (Scientific Editor, Department of Genetics, University Medical Center Groningen) for language editing, and Ren Mao for his kind suggestions on figure styling and structural improvements (Department of Gastroenterology, The First Affiliated Hospital of Sun Yat-Sen University, Sun Yat-Sen University, Guangzhou, Guangdong, China).

## Data availability

The datasets used and/or analyzed for the current study are available from the corresponding author on reasonable request. The data for the Groningen 1000IBD cohort can be requested with the accession number EGAS00001002702 (IDs: EGAD00001003991, EGAD00001008214, and EGAD00001008215).

## Code availability

All analytic code used for this study can be found at the following link: https://github.com/GRONINGEN-MICROBIOME-CENTRE/Groningen-Microbiome/tree/master/Projects/IBD_biopsy_project.

## Authors’ contributions

Conceptualization: RKW. Investigation: SH, ARB, RG, BHJ, RM, AB, IJH, JRB, HJMH, AVV, LMS and RKW. Methodology: SH, ARB, RG, BHJ, RM, IJH, JRB, HJMH, AVV,

LMS and RKW. Funding acquisition: RKW. Supervision: EAMF, AVV, LMS and RKW. Writing - original manuscript: SH, ARB and LMS. Writing - review and editing: all authors.

## Supplementary Methods

### Polymerase chain reaction (PCR), DNA clean-up, and MiSeq library preparation for mucosal 16 microbiota characterization

The PCR procedure consisted of the following conditions: an initial cycle of 94°C for 3 min followed by 32 cycles of 94°C for 45 sec, 50°C for 60 sec and 72°C for 90 sec, with a final extension of 72°C for 10 min. Agarose gel electrophoresis confirmed the presence of the PCR product (band at ∼465 bp) in successfully amplified samples.

Subsequently, DNA samples were thoroughly cleaned by mixing the remainder of the PCR product with 25 μL Agencourt AMPure XP beads (Beckman Coulter, Brea, California, USA) followed by an incubation of 5 min at room temperature. Beads were separated from the mixture by placing the samples within a magnetic bead separator for 2 min. After discarding the cleared solution, beads were washed twice by resuspending them in 200 μL fresh 80% ethanol, followed by an incubation of 30 sec in the magnetic bead separator, and again discarding the cleared solution. The pellet was dried for 15 min and resuspended in 52.5 μL 10 mM Tris HCl buffer (pH 8.5). Fifty (50) μL of this solution was subsequently brought into a new tube. DNA concentrations were measured using a Qubit^®^ 2.0 fluorometer (Thermo-Fisher Scientific, Waltham, Massachusetts, USA). To ensure similar library representations across samples, 2 nM dilutions of each sample were prepared accordingly. A library was created by pooling 5 μl of each diluted sample. Subsequently, 10 μL of the sample pool and 10 μL 0.2 M NaOH were mixed and incubated for 5 min to allow denaturation of the sample DNA.

980 μL of the HT1 buffer of the MiSeq 2x300 cartridge was then added to this mixture. Next, a denatured diluted PhiX solution was created by combining 2 μL 10 nM PhiX library with 3 μL 10 mM Tris HCl buffer (pH 8.5) with 0.1% Tween-20. 5 μL 0.2 M NaOH was added to this mixture and incubated for 5 min at room temperature. This 10 μL mixture was eventually mixed with 990 μL HT1 buffer. From the diluted sample pool, 150 μL was combined with 50 μL of the diluted PhiX solution, which was further diluted by the addition of 800 μL HT1 buffer. Finally, 600 μL of the prepared library solution was loaded into the sample loading reservoir of the 2x300 MiSeq cartridge for 16S rRNA amplicons sequencing (MiSeq Benchtop Sequencer, Illumina, San Diego, California, USA). Samples with low DNA concentrations after clean-up (quality score < 0.9) were discarded by PANDAseq to increase quality of sequence read-outs.

**Supplementary Table S1.**
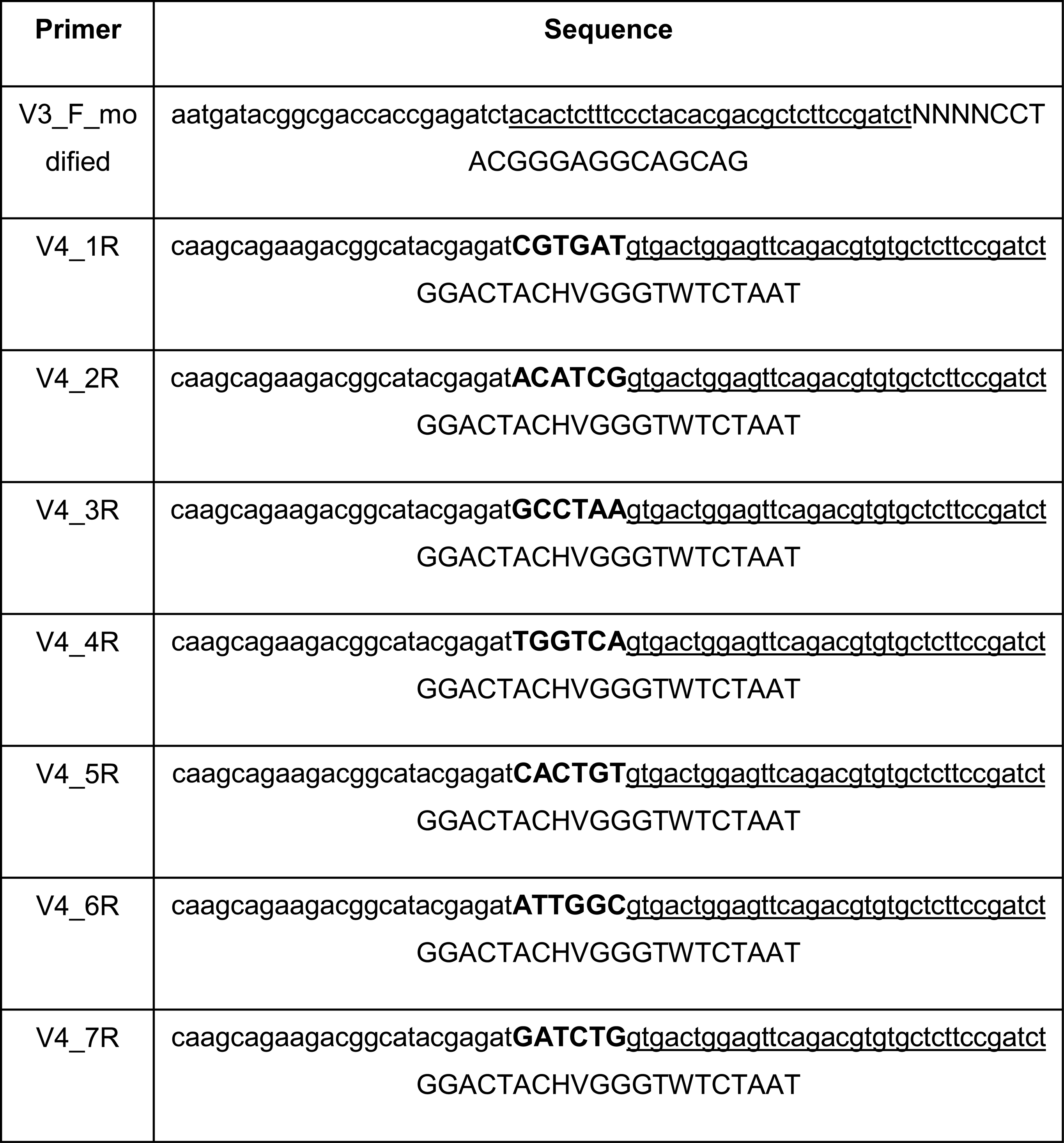
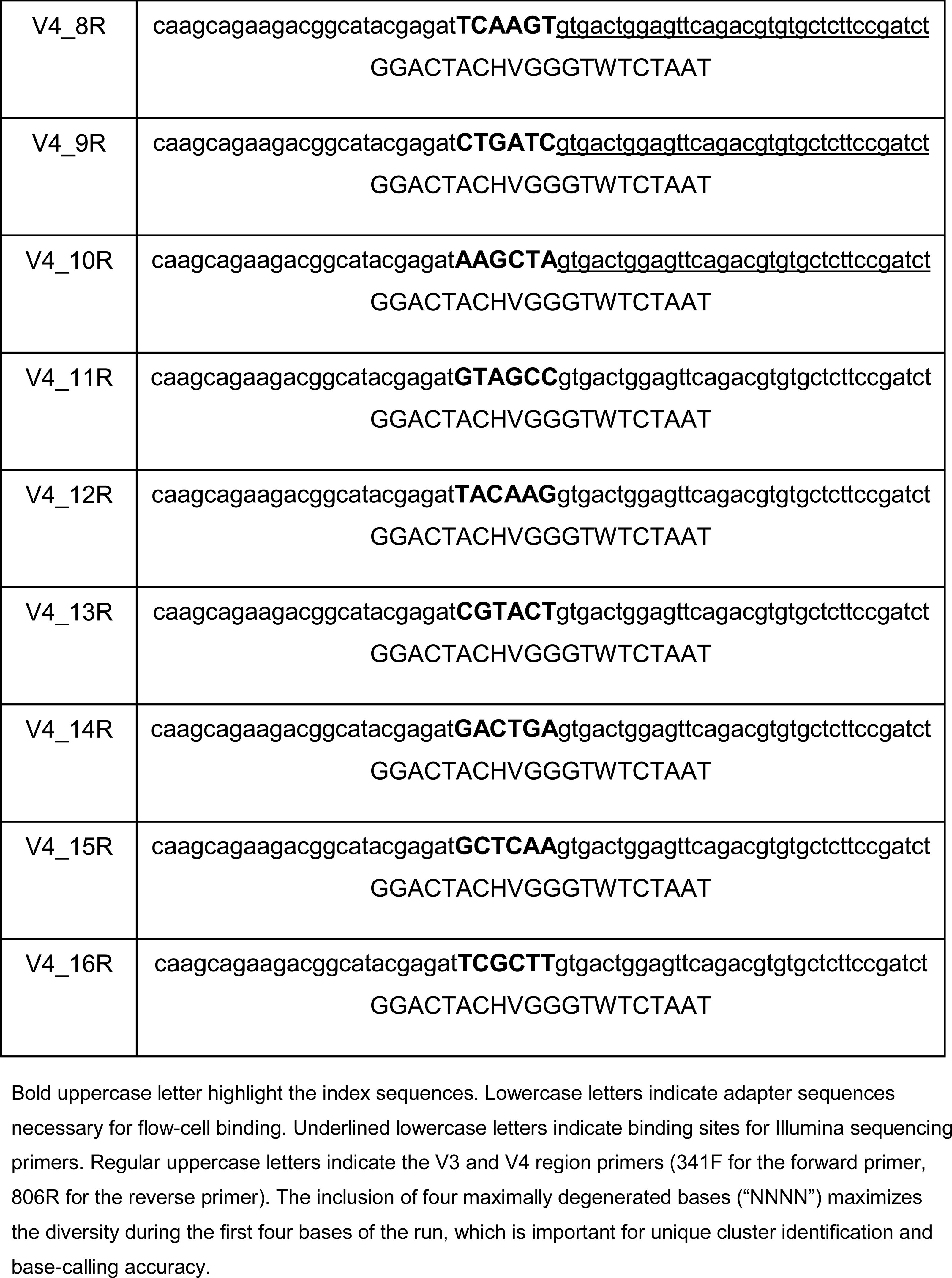
Nucleotide sequences of primers used for library construction for bacterial 16S rRNA gene (Illumina) sequencing.

## Supplementary Table Index

All Supplementary Tables have been uploaded separately for peer-review.

**Supplementary Table S1**. Differential gene expression analyses between non-inflamed and inflamed biopsies from ileal CD (group 1), colonic CD (group 2) and UC (group 3).

**Supplementary Table S2**. Differential gene expression analysis between inflamed biopsies from patients with CD (reference) and patients with UC.

**Supplementary Table S3**. Differential expression analysis of deconvoluted cell types between inflamed colonic biopsies of patients with CD (reference) and patients with UC.

**Supplementary Table S4**. Relative abundances of mucosal bacterial groups in different groups (CD, UC and non-IBD controls) and biopsy locations (ileum or colon).

**Supplementary Table S5**. Comparison of relative abundances of bacterial groups between non-IBD controls and ileal CD (group 1), colonic CD (group 2) and UC (group 3).

**Supplementary Table S6**. Hierarchical analysis performed using an end-to-end statistical algorithm (HAllA) demonstrating the main associations between mucosal bacterial groups and clinical phenotypes.

**Supplementary Table S7**. Genes and bacteria contained in component pair 1 from sparse-CCA analysis (FDR<0.05).

**Supplementary Table S8**. Pathway annotation of genes involved in component pair 1 from sparse-CCA analysis (FDR<0.05).

**Supplementary Table S9**. Genes and bacteria contained in component pair 3 from sparse-CCA analysis (FDR<0.05).

**Supplementary Table S10**. Pathway annotation of genes involved in component pair 3 from sparse-CCA analysis (FDR<0.05).

**Supplementary Table S11**. Genes and bacteria contained in component pair 7 from sparse-CCA analysis (FDR<0.05).

**Supplementary Table S12**. Pathway annotation of genes involved in component pair 7 from sparse-CCA analysis (FDR<0.05).

**Supplementary Table S13**. Genes and bacteria contained in component pair 8 from sparse-CCA analysis (FDR<0.05).

**Supplementary Table S14**. Pathway annotation of genes involved in component pair 8 from sparse-CCA analysis (FDR<0.05).

**Supplementary Table S15**. Genes and bacteria contained in component pair 5 from sparse-CCA analysis (FDR<0.05).

**Supplementary Table S16**. Pathway annotation of genes involved in component pair 5 from sparse-CCA analysis (FDR<0.05).

**Supplementary Table S17**. Genes and bacteria contained in component pair 9 from sparse-CCA analysis (FDR<0.05).

**Supplementary Table S18**. Pathway annotation of genes involved in component pair 9 from sparse-CCA analysis (FDR<0.05).

**Supplementary Table S19**. Individual pairwise gene-bacteria associations.

**Supplementary Table S20**. Genes and bacteria associated with fibrostenotic CD/Montreal B2 (reference: Montreal B1).

**Supplementary Table S21**. Microbiota-associated gene clusters in patients with non- stricturing, non-penetrating disease (Montreal B1) and fibrostenotic CD (Montreal B2) including microbiota-associated pathway annotation and cluster comparisons.

**Supplementary Table S22**. Genes and bacteria associated with TNF-α-antagonists use (reference: non-users).

**Supplementary Table S23**. Microbiota-associated gene clusters in patients not using and using TNF-α-antagonists including microbiota-associated pathway annotation and cluster comparisons.

**Supplementary Table S24**. Individual pairwise gene-bacteria associations and their interaction with the degree of mucosal dysbiosis (Lloyd-Price et al., Nature 2019).

**Supplementary Table S25**. Mucosal microbiota and other phenotypic factors explaining variation in mucosal cell type enrichment in patients with IBD.

## Extended Data Figures

**Extended Data Fig. S1.**
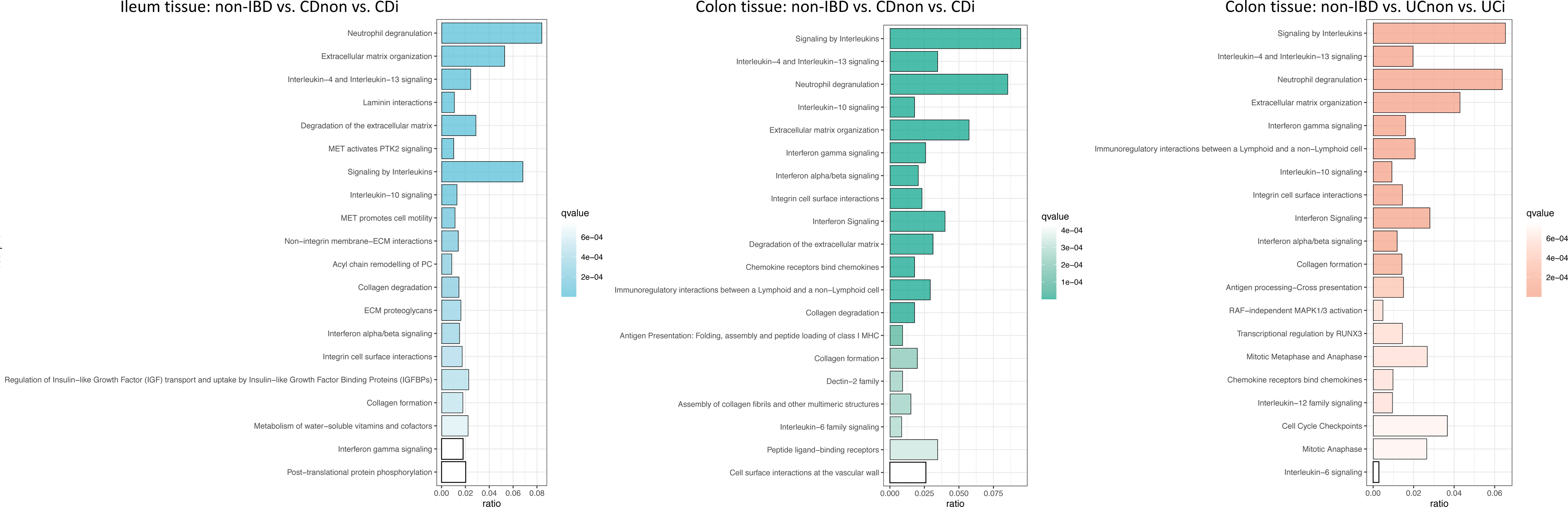
Analysis of pathways associated with each comparative gene expression analysis. The main pathways associated with inflamed ileal tissue in patients with CD (blue) include neutrophil degranulation, extracellular matrix (ECM) organization and IL-4/IL-13-signaling. Similar pathways were overexpressed in inflamed colonic tissue from patients with CD (green), but with a more prominent contribution from interleukin signaling pathways. Interleukin signaling pathways were also dominantly expressed in inflamed colonic tissue from patients with UC (orange), with other pathways expressed including neutrophil degranulation, ECM pathways, interferon gamma signaling and immunoregulatory interactions between lymphoid and non-lymphoid cells. Pathways were annotated using the Reactome pathway database. Abbreviations: CDi, inflamed tissue from patients with Crohn’s disease; CD-non, non-inflamed tissue from patients with Crohn’s disease; UCi, inflamed tissue from patients with ulcerative colitis; UC-non, non-inflamed tissue from patients with ulcerative colitis.

**Extended Data Fig. S2.**
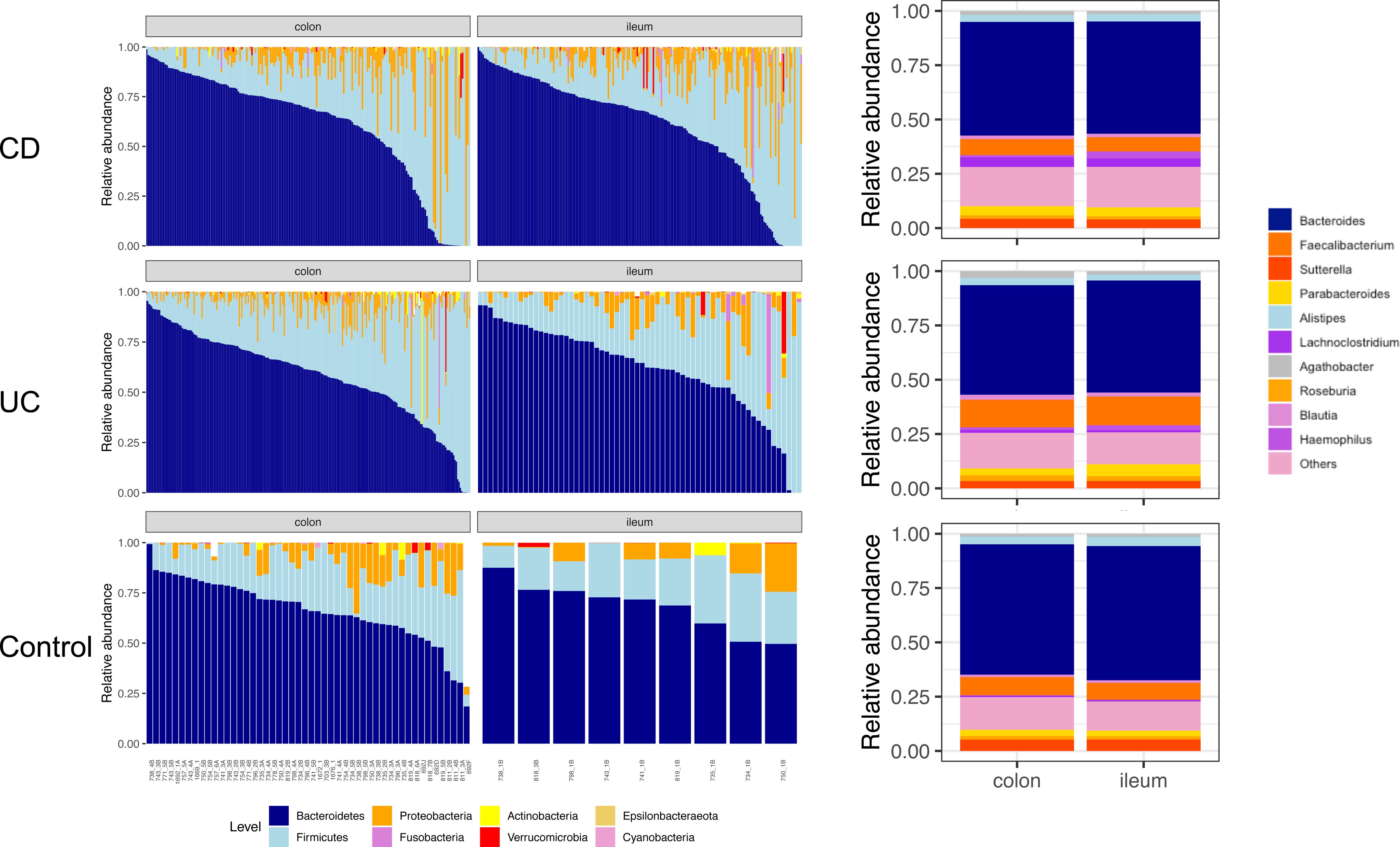
Mucosal 16S rRNA gene sequencing characterization demonstrates distinct compositional differences in relative abundances on. (**A**) bacterial phylum level and (**B**) bacterial genus level. Abbreviations: CD, Crohn’s disease; UC, ulcerative colitis.

**Extended Data Fig. S3.**
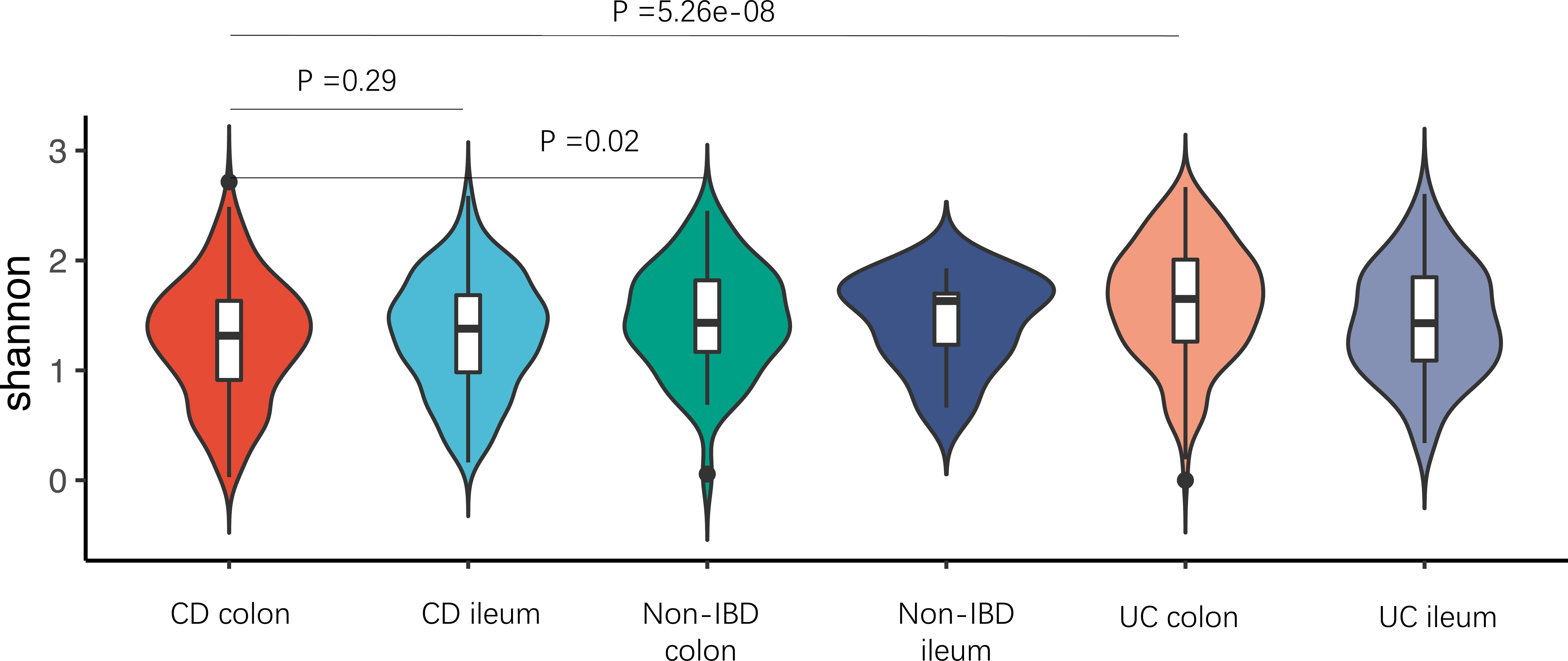
Microbial alpha-diversity (Shannon index) is significantly lower in colonic biopsies from patients with CD compared to colonic biopsies derived from patients with UC or controls. This indicates that this difference is not solely attributable to ileal biopsies from patients with CD.

**Extended Data Fig. S4.**
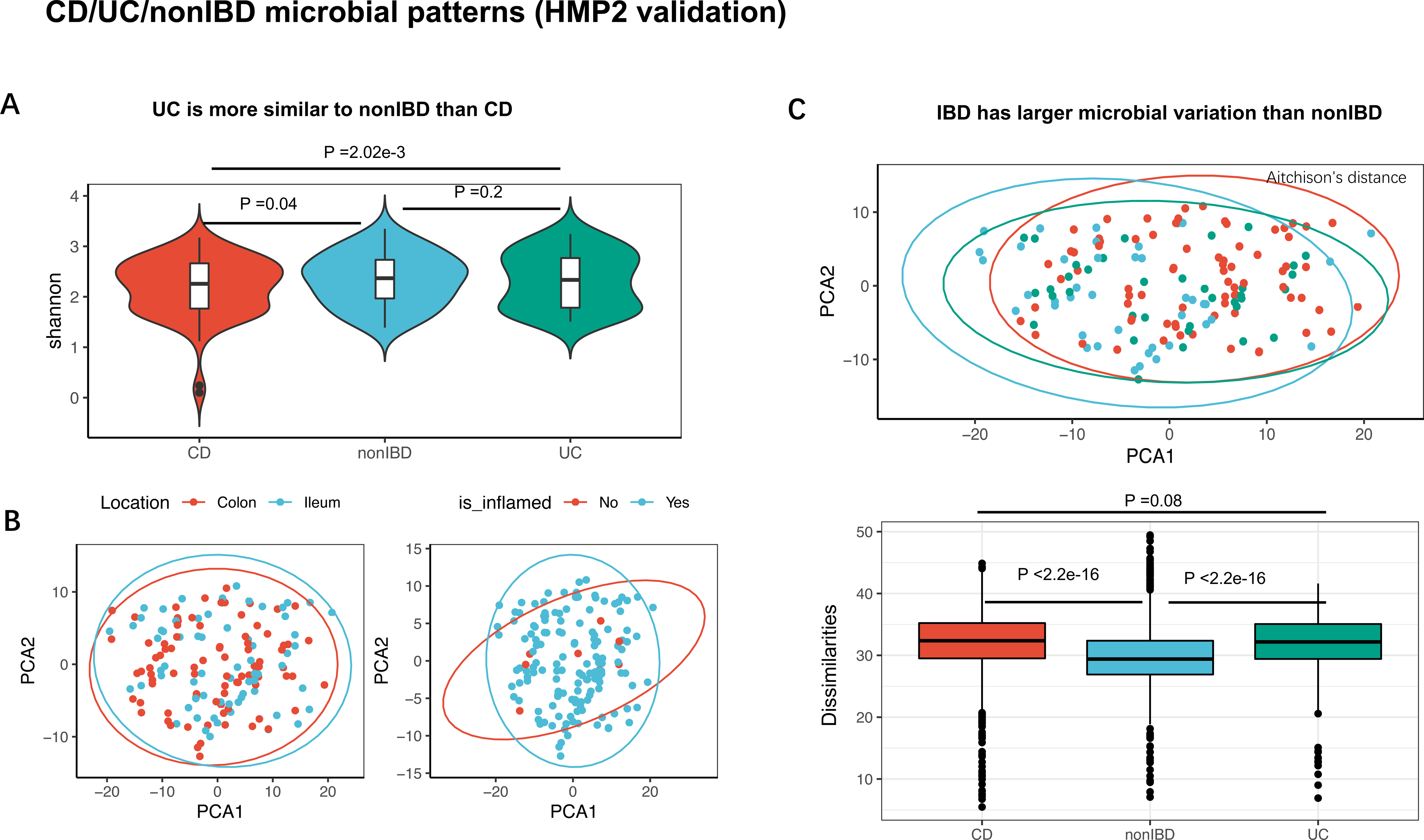
Replication of overall mucosal microbiota characterization in patients with IBD and non-IBD controls. Replication was performed in data derived from the HMP2 cohort study [13]. **a**, Microbial alpha-diversity (Shannon index) was lowest in ptaients with CD (n=85) compared to patients with UC (n=46) and non-IBD controls (n=45). **b**, PCA plots based on Aitchison’s distances and stratified by tissue location and inflammatory status (colors as in **a**). **c**, PCA plot showing microbial dissimilarity (Aitchison’s distances) in CD, UC and non-IBD controls. **d**, Microbial dissimilarity is highest in samples from patients with CD, followed by patients with UC and non-IBD controls. CD, Crohn’s disease; PCA, principal component analysis; UC, ulcerative colitis.

**Extended Data Fig. S5.**
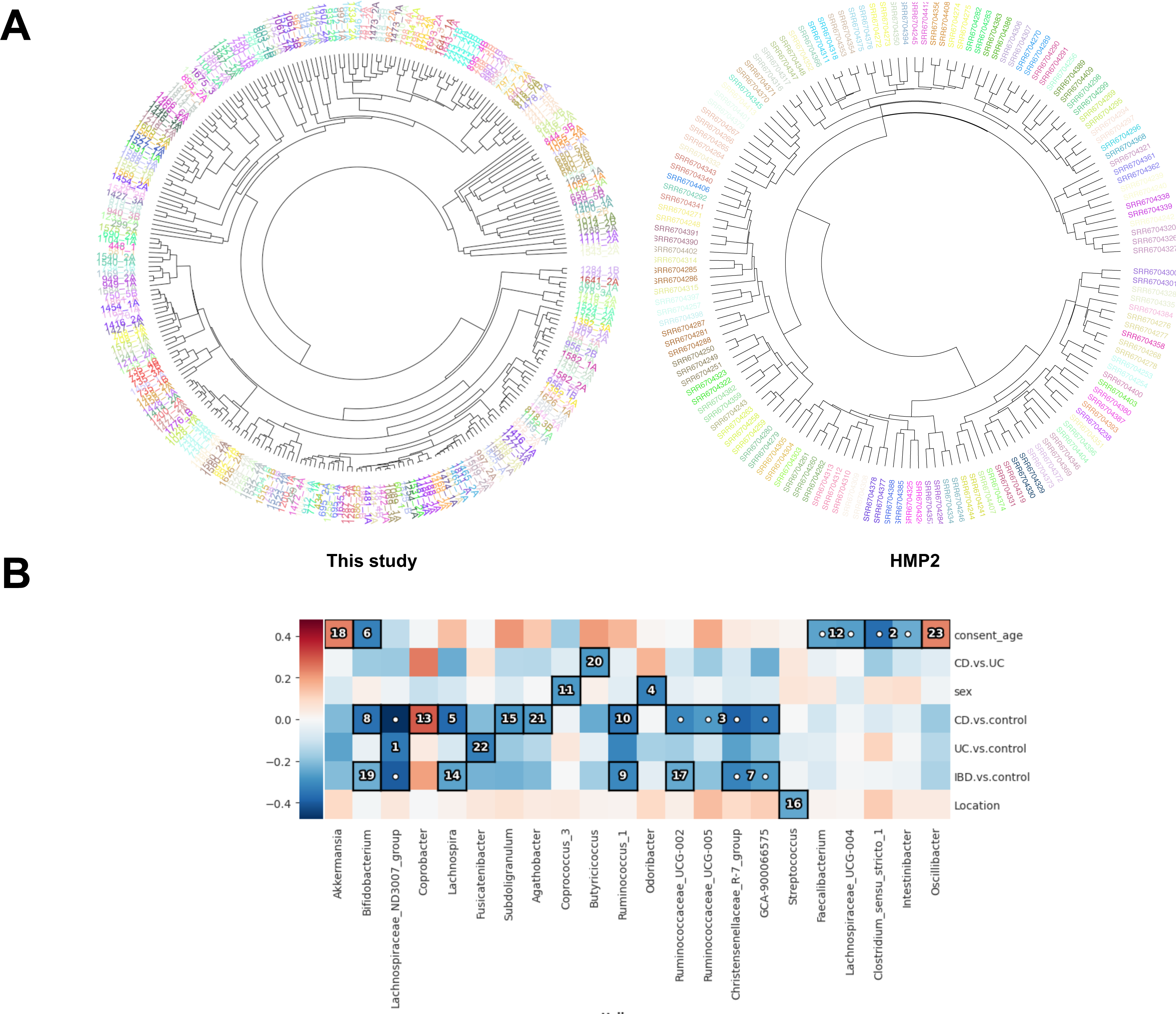
Composition of the mucosal microbiota is highly personalized and influenced by disease parameters and clinical factors in patients with IBD and controls. (**A**) Hierarchical clustering analysis demonstrating that tissue samples from the same individual (paired samples) clearly cluster together (colors indicate unique individuals). (**B**) Hierarchical analysis performed using an end-to-end statistical algorithm (HAllA) showing the main phenotypic factors that correlate with intestinal mucosal microbiota composition. Heatmap color palette indicates normalized mutual information. Numbers and dots in cells identify the significant pairs o features (phenotypic factors vs. bacterial taxa) in patients with IBD and controls. Abbreviations: BMI, body-mass index; CD, Crohn’s disease; UC, ulcerative colitis.

**Extended Data Fig. S6.**
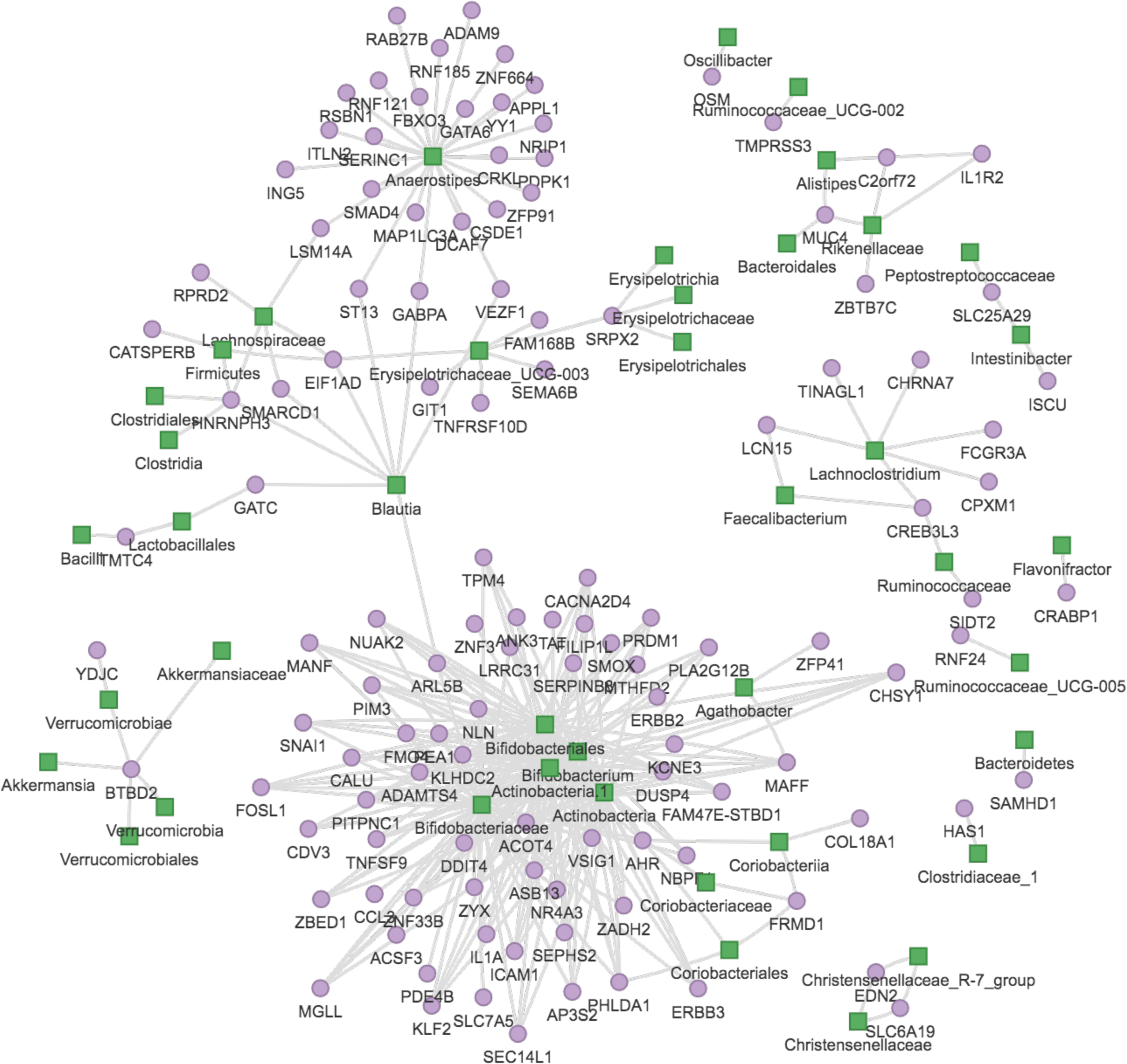
Network graph displaying significant individual gene–bacteria interactions. Green squares indicate bacterial groups. Purple dots indicate host gene expression. Each connecting line indicates statistically significant gene–bacteria associations after adjustment for age, sex, batch, medication use, tissue inflammatory status and tissue location. Most individual gene–bacteria associations (94%) overlap with the results from the sparse-CCA analysis (Figure 4).

**Extended Data Fig. S7.**
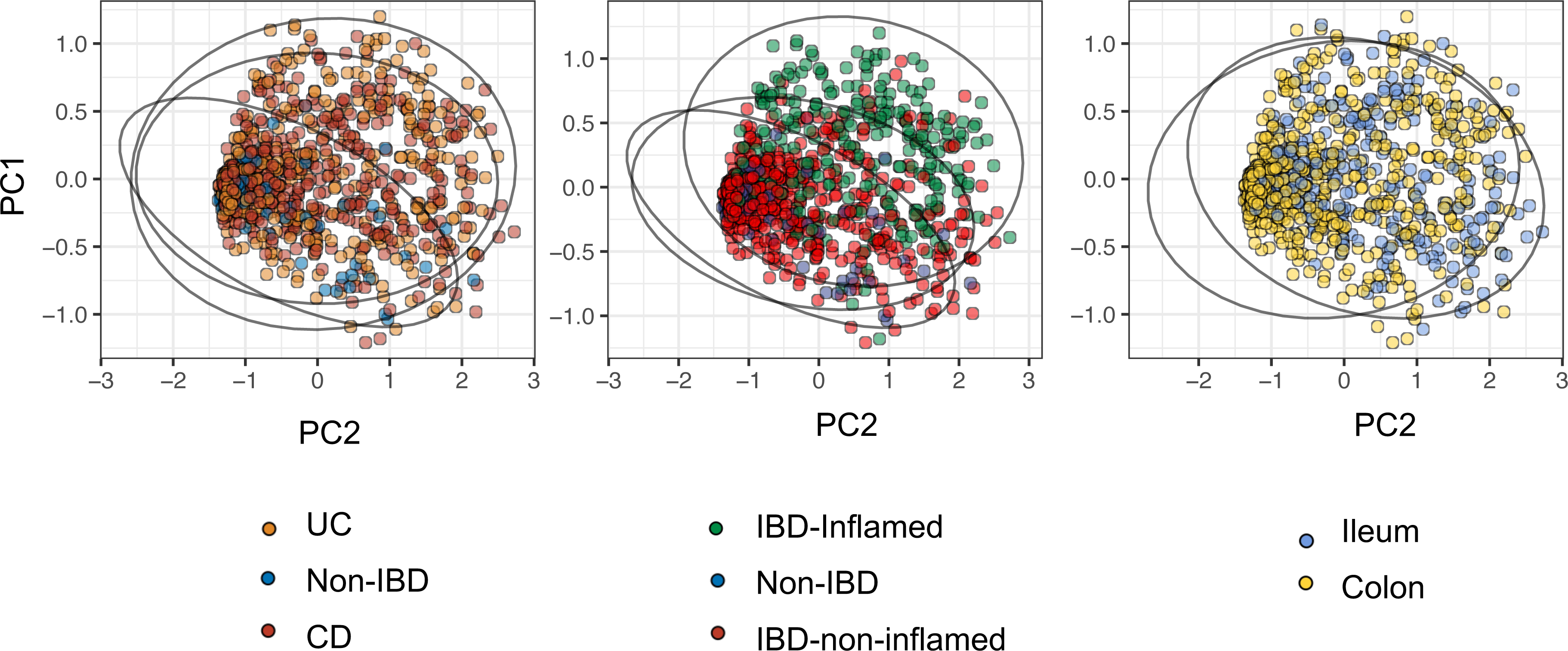
**Principal component analysis (PCA) plots demonstrating variation in cell type–enrichment labeled by diagnosis, biopsy inflammatory status and intestinal location**. Each dot represents one tissue sample. <u>Left</u>: Patients with IBD, both CD and UC, show significantly different intestinal cell type composition compared to controls. <u>Middle</u>: Tissue inflammatory status induces shifts in cell type composition, showing differences between non-inflamed IBD tissue vs. control tissue and inflamed IBD tissue vs. control tissue. <u>Right</u>: Tissue location (ileum vs. colon) also demonstrates distinct variation in cell type composition.

## Supplementary Results

### Box 1. Individual mucosal gene–bacteria associations and their potential biological implications in IBD

#### Mucosal bifidobacteria positively associate with aryl hydrocarbon receptor (AHR) and ABC-transporter (ABCC1) expression levels

The positive association between *AHR* expression and bifidobacteria could be explained by the fact that *Bifidobacterium* spp. can produce aromatic lactic acids such as indole-3-lactic acid (out of aromatic amino acids like tryptophan) via aromatic lactate dehydrogenase, which in turn activates the host aryl hydrocarbon receptor [1, 2]. Activation of the aryl hydrocarbon receptor, a crucial regulator of intestinal homeostasis and immune responses, leads to a reduction of inflammation in intestinal epithelial cells [3] and confers immunoprotective effects [4].

Another intriguing observation is the positive association between bifidobacteria and host expression of the *ABCC1* gene. *ABCC1* is a member of the ATP-binding cassette transporters (ABC transporters, and also known as multidrug resistance-associated protein 1, MRP1) that has multiple physiological functions, but it may also confer pathophysiological sequelae, especially in the context of cancer [5]. Under physiological circumstances, it detoxifies endogenously generated toxic substances (as well as xenobiotics), protects against oxidative stress, transports leukotrienes and lipids and may facilitate the cellular export and body distribution of vitamin B_12_ [6]. Interestingly, several *Bifidobacterium* species (e.g. *B. animalis*, *B. longum* and *B. infantis*) can synthesize vitamin B_12_, which is subsequently absorbed in the large intestine via unknown mechanisms [7–9].

#### Mucosal bifidobacteria associate with FOSL1, a subunit of the AP-1 transcription factor

Associations between mucosal *Bifidobacterium* bacteria and expression of *FOSL1* genes were amongst the top significant individual gene–bacteria interactions. Fos- related antigen 1 (FRA1), encoded by *FOSL1*, is a subunit of the activator protein 1 (AP-1) transcription factor. In the intestine, the AP-1 transcription factor is commonly activated in response to inflammatory stimuli and has been implicated in IBD [10]. More specifically, an interaction may exist between AP-1 activity and the glucocorticoid receptor, which may be part of the anti-inflammatory effects of steroid treatment [11]. In steroid-resistant patients with CD, AP-1 activation was primarily observed in the nuclei of intestinal epithelial cells, whereas this activation was restricted to lamina propria macrophages in steroid-sensitive patients [10]. This suggests a differing cellular activation pattern of AP-1 activation in steroid-resistant patients where the expression of this transcription factor may interfere with the activity of the glucocorticoid response. In an experimental study in which pregnant mice were supplemented with butyrate, FOS genes, including *Fosl1*, were observed to be downregulated in the colon and associated with protection against experimentally-induced colitis [12]. Although there are currently no reports of potential immune-modulating effects for Fosl1, it has 85% homology with Fosl2, another AP-1 transcription factor. A recent study demonstrated that Fosl2 is important in T-reg development and control of autoimmunity. Interestingly, several GWASs have reported associations of a SNP located in the promoter region of *FOSL2* with IBD [13–15], and the presence of this SNP was also shown to correlate with *FOSL2* expression in blood cells of patients with IBD [16]. In the context of T-regs, *FOSL2* also appears to be important as it is a determinant of a highly suppressive subpopulation of T-regs in humans that are particularly enriched in the lamina propria of patients with CD, supporting wound healing in the intestinal mucosa [17]. Although speculative, bifidobacteria, as well as their metabolites such as butyrate, may potentially confer immune-modulating properties via interaction with *FOSL1* expression.

#### Mucosal bifidobacteria positively associate with Krüppel-like factor 2 (KLF2) expression

Krüppel-like factor 2 (encoded by *KLF2*) is a negative regulator of intestinal inflammation, and its expression is found to be reduced in patients with IBD [18]. *KLF2* also negatively regulates differentiation of adipocytes and strongly inhibits PPAR-γ expression, which prevents differentiation of preadipocytes into adipocytes and thereby prevents adipogenesis [19]. *KLF2* also plays an important role in endothelial physiology, where it may act as a molecular switch by regulating endothelial cell function in inflammatory disease states [20]. Interestingly, *KLF2* modifies the trafficking of T-regs, as increased *KLF2* expression in T-regs promotes the induction of peripheral immunological tolerance, whereas, in the absence of its expression, T-regs are unable to effectively migrate to secondary lymphoid tissues [21]. Indeed,, it was demonstrated in mouse experiments that mice developed IBD in the presence of *KLF2*-deficient T- regs, which were unable to prevent colitis by disrupted co-trafficking of effector and regulatory T cells. In light of these considerations, mucosal bifidobacteria may confer beneficial immune-modulating properties by upregulating *KLF2* expression, thereby stimulating T-reg migration and contributing to immunological self-tolerance in the context of IBD.

#### Mucosal Anaerostipes bacteria positively associate with host SMAD4 expression

*Anaerostipes*, which belong to the *Lachnospiraceae* family, are anaerobic bacteria that are well-known butyrate-producers. Butyrate serves as the primary energy source for colonic epithelial cells and is characterized by anti-inflammatory and anti-carcinogenic properties. *SMAD4* is an important intracellular effector of the TGF-β superfamily of proteins. These proteins have important functions in alleviating intestinal inflammation and maintenance of gut mucosal homeostasis. Haploinsufficiency of *SMAD4* in mice and humans has been associated with an increased susceptibility to colonic inflammation [22]. In patients with CD, reduced epithelial protein levels of SMAD4 were observed that was associated with disease activity, indicating defective mucosal TGF-β signaling during active intestinal inflammation. In an experimental animal study, mice with an epithelial deletion of *Smad4* presented with macroscopic invasive adenocarcinoma of the distal colon and rectum 3 months after DSS-induced colitis [23]. Indeed, *SMAD4* mutations in humans are linked to juvenile polyposis syndrome and associated with poor disease outcome in several types of cancer [24–27]. Using RNA-seq analysis, a strong inflammatory expression profile was observed after *SMAD4* deletion, with expression of various inflammatory cytokines and chemokines, including CCL20. In addition, it was demonstrated that CCL20 could be repressed by *SMAD4* in colonic epithelial cells, proving that TGF-β signaling could block the induction of CCL20 expression to protect against the development of colitis-associated cancer.

In an experimental study involving human hepatic stellate cells, butyrate was demonstrated to be protective against diet-induced nonalcoholic steatohepatitis and liver fibrosis via suppression of TGF-β signaling pathways in which SMAD proteins are involved. Although butyrate mainly showed antifibrotic effects via reduction of non-canonical TGF-β signaling cascades, there was also a significant increase in the expression of SMAD4 with the addition of butyrate on top of TGF-β treatment [28]. We found *Anaerostipes* bacteria to also be strongly associated with expression of *ZNF644*, a zinc finger protein that is positively regulated by intracellular zinc concentrations.

Depletion of intracellular zinc levels, or even zinc deficiency, may have destabilizing effects on SMAD proteins and thereby impair the TGF-β signaling pathway [29].

#### Mucosal Verrucomicrobia bacteria inversely associate with expression of the IBD susceptibility gene YDJC

We observed significant inverse associations between Verrucomicrobia bacteria, of which *Akkermansia muciniphila* is a well-known member, and the expression of the *YDJC* gene, which encodes for the YdjC chitooligosaccharide deacetylase homolog (YdjC) protein. This gene has been identified as a shared susceptibility gene for CD, UC and psoriasis [13,30,31]. *YDJC* was originally identified as a celiac disease–associated susceptibility locus, but some SNPs were also associated with CD as well as with pediatric-onset CD [32]. YdjC catalyzes the deacetylation of acetylated carbohydrates, an important reaction in the degradation of oligosaccharides [33]. *YDJC* expression has been associated with tumor progression in studies of lung cancer [34, 35]. The observed inverse association between *Akkermansia* and *YDJC* expression may suggest a potential protective role of *Akkermansia*, as decreased *YDJC* expression may mitigate its pro-carcinogenic effects. Despite the association between *YDJC* and the susceptibility to IBD on a genetic level, its precise functional role remains largely unknown [32].

#### Mucosal Alistipes bacteria positively associate with MUC4 expression

The bacterial genus *Alistipes*, belonging to family *Rikenellaceae* and phylum Bacteroidetes, is a recently discovered bacterial species, of which many have been isolated from the human gut microbiome. The role of *Alistipes* in health and disease is still unclear. Some evidence indicates that it may confer protective effects to the host, but other studies report pathogenic effects, e.g. in colorectal cancer development. A key factor believed to determine the relative abundance of *Alistipes* is the dysbiotic state of the gut microbiome [36]. In IBD, there is also conflicting data about the pathogenicity of *Alistipes* species. *Alistipes finegoldii* has been demonstrated to exert anti-inflammatory effects in experimental models of colitis [37]. Likewise, another study found an increased abundance of *Alistipes* in *NOD2*-knockout mice that had less severe (TNBS-induced) colitis compared to wild-type mice [38]. It has also been reported that *Alistipes* abundance could increase after taking probiotic supplements, which in turn may protect against hepatocellular cancer growth in an experimental setting [39]. However, metagenomic studies have shown that *Alistipes* abundances were increased in mouse models of spontaneous CD-like ileitis terminalis as compared to wild-type mice, suggesting that *Alistipes* species may also play a pathogenic role by eliciting segmental ileitis [40, 41].

*MUC4* encodes for mucin 4, a protein found in the glycocalyx present on the intestinal epithelium. Deletion or knockouts of *Muc4* have demonstrated protective effects in mouse models, as shown by lower levels of proinflammatory factors and resistance against DSS-induced colitis. It is still unclear how this protective mechanism of *MUC4* deletion works, but it has been hypothesized that it may trigger the concomitant upregulation of other mucin proteins (e.g. *MUC*1-3) as these genes have been observed to be highly expressed in *Muc4*-knockout mice with DSS-induced colitis [42, 43]. Based on this, we speculate that the positive association between *Alistipes* abundance and *MUC4* expression may imply a potential pathogenic role of *Alistipes* in the context of IBD-associated dysbiosis. However, in our data, we did not observe a significant interaction via dysbiotic status between *Alistipes* abundance and *MUC4* expression.

#### Mucosal Oscillibacter bacteria positively associate with OSM expression

*Oscillibacter*-like bacteria, which include *Oscillibacter* and *Oscillospira*, are commonly detected in human gut microbial communities, although their exact physiological role is not fully understood. Previously, it was reported that *Oscillibacter* may be a potentially important bacterium in mediating high fat diet–induced intestinal dysfunction, which was supported by a negative association between *Oscillibacter* and intestinal barrier function parameters [44]. Similarly, the abundance of *Oscillibacter* has been reported as a key bacterial group associated with colitis development in DSS-induced colitis in mice and with prenatal stress in rodents [45, 46]. However, a recent study linking gut microbiota profiles to sulfur metabolism in patients with CD demonstrated that *Oscillibacter* abundance was enriched in patients with inactive compared to active disease but diminished in patients with IBD compared to controls [47, 48]. Thus, similar to *Bacteroides* and *Alistipes*, the exact functional role of *Oscillibacter* in the context of IBD remains elusive, but it will likely depend on gut microbial dysbiosis and the intestinal (inflammatory) environment. The *OSM* gene encodes for the oncostatin M protein, a well-known inflammatory mediator in IBD that drives intestinal inflammation, mainly *via* activation of JAK-STAT and PI3K-Akt pathways [49]. Besides induction of other inflammatory events, it primarily triggers the production of various cytokines, chemokines and adhesion molecules that contribute to intestinal inflammation [50]. In addition, OSM is a marker for non-responsiveness to TNF-α-antagonists in patients with IBD [51]. Considering these findings, the positive association between *OSM* expression and *Oscillibacter* abundance we observe supports a potentially pathogenic role for this bacterial species in IBD.

#### Mucosal Blautia bacteria associate with host ST13 expression levels

Hsc70-interacting protein, encoded by the *ST13* gene, mediates the assembly of the human glucocorticoid receptor, which requires involvement of intracellular chaperone proteins such as heat shock proteins HSP70 and HSP90 [52]. Reduced expression of ST13 has been observed in patients with colorectal cancer, suggesting that ST13 may constitute a candidate tumor-suppressor gene [53, 54]. The positive association we observe between mucosal *Blautia* abundance and *ST13* gene expression may therefore point to a protective anti-carcinogenic role for *Blautia* in the intestines.

Supplementary References to Box 1

## Box 2. Miscellaneous component pairs from sparse-CCA analysis

The microbial part of the fifth pair of components (component pair 5, *P*=2.87 x10^-8^, FDR<0.05) was formed by *Christensenellaceae*, *Ruminococcaceae*, *Lachnospiraceae* (NK4A136 group), Coriobacteria and the genera *Coprococcus* and *Ruminoclostridium*, which are all inversely associated with pathways representing SLC-mediated transmembrane transport (e.g. transport of bile acids and organic acids, metal ions and amine compounds) as well as biological oxidation and fat metabolism pathways including arachidonic acid metabolism and (glycero)phospholipid biosynthesis (**Supplementary Tables S15-S16**).

In the sixth pair of components (component pair 9, *P*=9.65 x10^-7^, FDR<0.05), the microbial component was primarily composed of bifidobacteria (i.e. order Bifidobacteriales, family *Bifidobacteriaceae* and genus *Bifidobacterium*), which were inversely associated with pathways representing phospholipid synthesis (e.g. phosphatidic acid synthesis) and NR1H2/NR1H3 or liver X receptor (LXR)-mediated signaling (**Supplementary Tables S17-S18**). NR1H3 (LXR-α) and NR1H2 (LXR-β) are ligand-activated transcription factors stimulated by endogenously produced oxysterols, which are in turn produced by oxidation of cholesterol, enzymatic reactions or alimentary processes [1]. Under physiological conditions, oxysterols are formed proportional to the cellular cholesterol content and thereby stimulate LXRs (acting as cholesterol sensors) to alter gene expression and activate protective mechanisms to prevent cholesterol overload in the cell. This occurs via inhibition of intestinal cholesterol absorption, activation of cholesterol efflux from cells to HDL (via ABCA1 and ABCG1 transporters) and activation of the hepatic conversion of cholesterol to bile acids and stimulation of biliary cholesterol and bile acid excretion. In addition, LXR-agonists enhance *de novo* synthesis of fatty acids by stimulating the expression of the lipogenic transcription factor SREBP-1c, which may result in elevated plasma triglycerides and hepatic steatosis. LXRs are also involved in modulation of innate and adaptive immune responses and regulate diverse aspects of inflammatory gene expression in macrophages. The ability of LXRs to coordinate metabolic and immune response constitutes an attractive therapeutic target for treatment of IBD.

